# Deep generation of personalized connectomes based on individual attributes

**DOI:** 10.1101/2025.02.12.637986

**Authors:** Yuanzhe Liu, Caio Seguin, Sina Mansour L., Ye Ella Tian, Maria A. Di Biase, Andrew Zalesky

## Abstract

An individual’s connectome is unique. Interindividual variation in connectome architecture associates with disease status, cognition, lifestyle factors, and other personal attributes. While models to predict personal attributes from a person’s connectome are abundant, the inverse task— inferring connectome architecture from an individual’s personal profile—has not been widely studied. Here, we introduce a deep model to generate a person’s entire connectome exclusively based on their age, sex, body phenotypes, cognition, and lifestyle factors. Using the richly phenotyped UK Biobank connectome cohort (N=8,086), we demonstrate that our model can generate network architectures that closely recapitulate connectomes mapped empirically using diffusion MRI and tractography. We find that age, sex, and body phenotypes exert the strongest influence on the connectome generation process, with an impact approximately four times greater than that of cognition and lifestyle factors. Regional differences in the importance of measures were observed, including an increased importance of cognition in the association cortex relative to the visual system. We further show that generated connectomes can improve the training of machine learning models and reduce their predictive errors. Our work demonstrates the feasibility of inferring brain connectivity from an individual’s personal data and enables future applications of connectome generation such as data augmentation and anonymous data sharing.

## Introduction

Understanding the structure and function of human brain networks remains a fundamental goal in neuroscience research. Advances in neuroimaging have enabled mapping of the human connectome (Hagmann, 2005; Sporns et al., 2005) at a variety of resolutions (Mansour L et al., 2021). These brain networks are complex, manifesting a cost-efficient topology that facilitates interregional communication and supports complex cognitive functions and behaviors (Bullmore & Sporns, 2012).

No two connectomes are the same and interindividual connectome variation associates with individual differences in demographics, developmental stages, health status, behaviors, and other factors. For example, structural connectomes exhibit higher modularity in early and late stages of life relative to young adulthood (Puxeddu et al., 2020), and network efficiency increases as individuals transition from childhood to adolescence (Feng et al., 2023). Sex differences in structural connectivity are also evident, with females showing stronger inter-hemispheric and inter-modular connectivity than males (Ingalhalikar et al., 2014; Tunç et al., 2016). In addition, connectome topology has been linked to cognitive performance (Akarca et al., 2021) and multiple neuropsychiatric conditions (Griffa et al., 2013; van den Heuvel & Sporns, 2019).

Building on these findings, numerous studies have developed models to predict an individual’s age, sex, and cognitive performance from their connectome (Kawahara et al., 2017; Kopetzky et al., 2024; Seguin et al., 2020; Yeung et al., 2023; Yeung et al., 2024). However, the inverse task— inferring connectomes from a individuals’ personal profile—remains largely unexplored. Solving this inverse problem can provide insights into how individual factors shape brain connectivity and paves the way for precision medicine through simulating the impact of an intervention or pharmacotherapy on an individual’s connectome. For instance, generating longitudinal connectomes for individuals enables prediction of personalized brain aging trajectories and the progression of a brain disorder, facilitating early and precise interventions. Modelling connectivity alterations in response to lifestyle changes may help identify behavioral modifications that can mitigate vulnerability to brain disorders.

Connectome generative models are not a new concept. Conventional connectome generative models are governed by a small number of wiring parameters that can be tuned to generate networks that match the properties of an individual’s empirically mapped connectome (Betzel et al., 2016; Bobyleva et al., 2024; Liu et al., 2023; Simpson et al., 2011; Song et al., 2014; Vértes et al., 2012). While interindividual variability in wiring parameters associates with age, sex, clinical diagnosis, and other individual measures (Betzel et al., 2016; Carozza et al., 2022; Vértes et al., 2012; Zhang et al., 2021), reported associations are typically weak and can vary depending on dataset and methodological choices. Moreover, current connectome generative models do not recapitulate node location (Liu et al., 2024), misplace brain hubs within the network (Oldham et al., 2021), and inaccurately reproduce long-range connections (Oldham et al., 2024).

In contrast, deep learning models can generate networks that better resemble empirically mapped connectomes (Liu et al., 2021). Early work used a graph autoencoder to learn the latent representation of connectivity, with extended functionality to generate connectomes based on a cognitive measure through Gibbs sampling (Liu et al., 2021). Following this framework, motion-aware deep generative models have been shown to reconstruct structural connectomes that more strongly associate with cognitive measures than raw connectivity (Zhang et al., 2024). However, debates persist regarding the use of deep models for connectome generation and prediction (Chen et al., 2024; Jamison et al., 2024; Smolders et al., 2024; Zalesky et al., 2024), with critics questioning whether these models truly capture interindividual connectome variability or if they merely learn to generate group-averaged connectomes. Despite these controversies, it remains unknown whether a person’s connectome *can* be generated solely based on their personal profile.

Here, we develop a generative model to parse an individual’s personal profile (i.e., age, sex, cognition, body phenotypes, and lifestyle factors) and generate their entire connectome based on this information. We then generate connectomes for individuals participating in the UK Biobank (Mansour L et al., 2023; Sudlow et al., 2015) and compare them to connectomes empirically mapped using diffusion MRI and tractography. We evaluate the extent to which our generated connectomes recapitulate interindividual variability and idiosyncrasies evident in the empirical connectomes. Our work provides a foundation for future applications of connectome generation, such as data augmentation, outlier detection, and anonymous data sharing.

## Results

### Connectome generation

We developed a deep model to generate structural connectomes for individuals based solely on their personal information. We used data from the UK Biobank (UKB), comprising a set of 194 individual measures grouped into 4 categories: age and sex, lifestyle factors, body phenotypes, and cognitive measures (Sudlow et al., 2015; Tian et al., 2023). Supplementary Table S1 catalogues all individual measures used as inputs to the generative process. These measures were selected based on established evidence linking them to structural connectivity (Feng et al., 2023; Ingalhalikar et al., 2014; Kawahara et al., 2017; Yeung et al., 2023). We considered a total of 8,086 UKB participants (age: 62.7 ± 7.3, 3,648 males) based on availability of personal data and structural connectomes, as well as absence of any self-reported and healthcare records documented clinical diagnosis. Our previously mapped connectomes were used for all experiments (Mansour L. et al., 2023). We specifically considered connectomes mapped using probabilistic tractography, weighted by streamline counts in the 200-node Schaefer parcellation delineated over Yeo’s 7 functional networks (Schaefer et al., 2018; Yeo et al., 2011). Structural connectomes in the Desikan-Killiany (DK) atlas were employed for replication of the results wherever applicable (see supplementary). Further details on data selection and preprocessing can be found in Methods.

Participants were randomly assigned into 80% training (*N* = 6,469, *age* = 62.7 ± 7.3, 2,935 males) and 20% test (*N* = 1,617, *age* = 62.4 ± 7.2, 713 males) sets. A generative model was trained to generate connectomes for unseen individuals in the test set using a set of 194 personal measures. All connectomes were represented in terms of a weighted connectivity matrix. We refer to connectomes generated by our model as *generated connectomes*, whereas connectomes mapped empirically with diffusion MRI and tractography are referred to as *empirical connectomes* (Fig. 1A).

**Figure 1.**
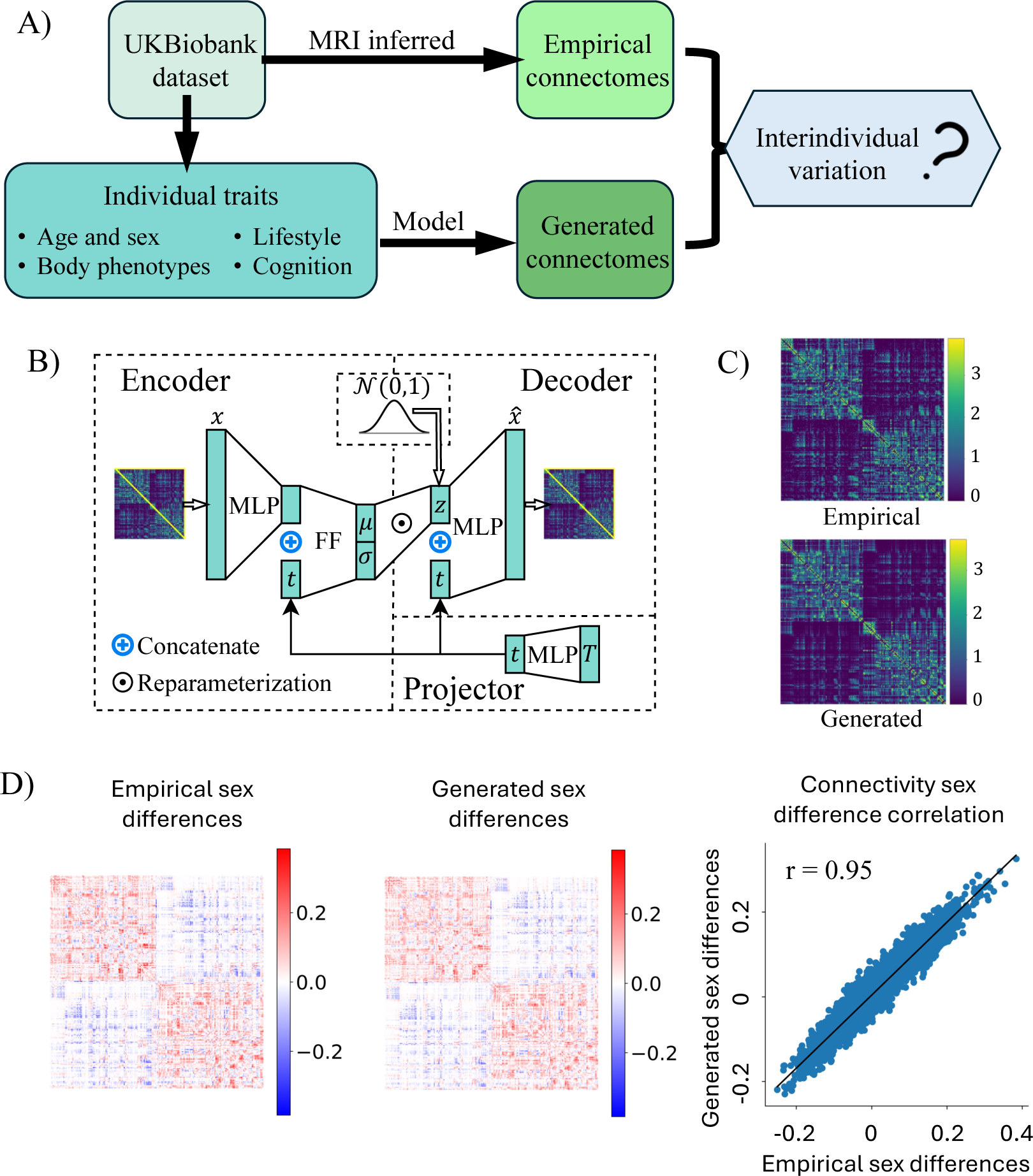
Overview of connectome generation and the generative model (cVAE) architecture. A) Model-generated connectomes (conditional on individual personal measures) are compared to empirical connectomes. B) Breakdown of cVAE architecture into the encoder, decoder, and projector. During training, latent representation *z* is sampled from the learned mean (μ) and log variance (σ) of latent space distributions using reparameterization. During generation, *z* is sampled from a standard normal distribution. Abbreviations: MLP – multilayer perceptron, FF: a single feedforward layer. C) The empirical and generated connectomes of an example subject in the test set. D) Generated connectomes resemble empirically observed sex differences in connectivity. Left visualizes the differences between male and female group average connectivity in the test set. Positive values indicate stronger connectivity in males than females. Center displays the sex differences in generated test set connectomes. The scatter plot on the right compares the empirical and generated sex differences in connectivity, with each data point representing a unique edge.

The generative model is a conditional variational autoencoder (cVAE) (Kingma, 2013) consisting of a profile projector, an encoder, and a decoder (Fig. 1B, see detailed architecture in Methods). The profile projector converts individual measures of continuous and binary variables *T* into a learnable hidden representation *t*, which is used as the conditional inputs to the encoder and the decoder. The encoder maps the distributions of latent representation *z* given the upper triangular elements of connectome *x* and the profile representation *t*, whereas the decoder reconstructs the connectivity edge weights 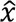 based on **z** and **t**. The model was trained on the training set to minimize the combined reconstruction and KL divergence loss with a Gaussian prior through stochastic gradient descent.

The trained model can generate connectomes conditional on an individual’s personal data (without knowing *x*, i.e., the structural connectivity) by sampling the latent representation *z* from a standard Gaussian distribution. Fig. 1C compares the empirical and generated connectomes of an example individual in the test set. Evaluating the trained model with the test set, we found that our generated and empirical connectomes were highly correlated within the same individuals (*r* = 0.92 ± 0.02). Between-group differences evident in the empirical data were also reflected in the generated networks (Fig. 1D, Fig. S1).

### Generated connectomes capture interindividual variation

We next quantified whether our generated connectomes preserved individual variability in connectome architecture. We used established identifiability and network measures (Fig. 2A and Methods) to quantify individual variability. Model-generated individual variability was benchmarked to a null model, as well as a state-of-the-art matching index (MI) model (Betzel et al., 2016) for connectome generation wherever appropriate (Methods).

**Figure 2.**
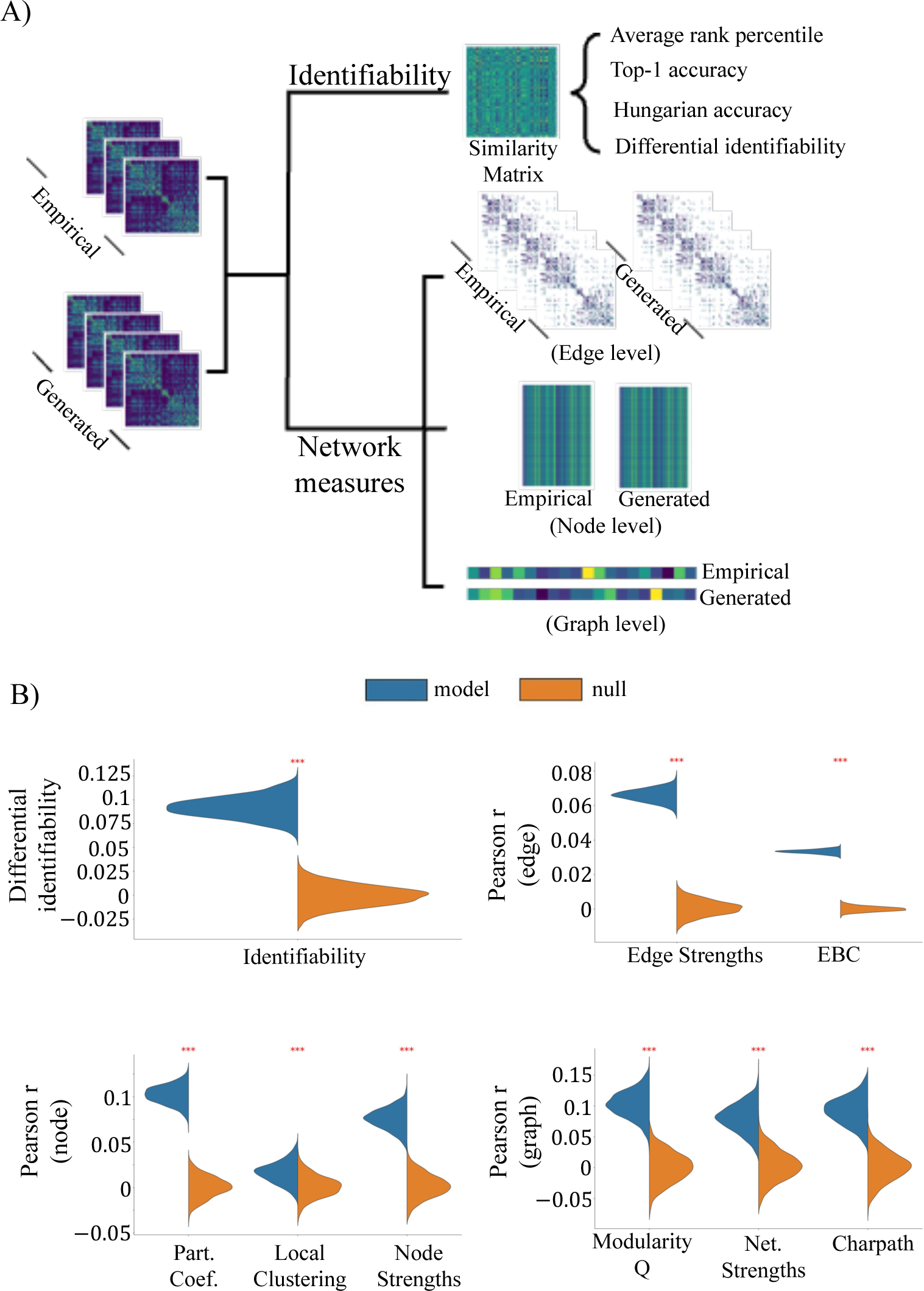
Evaluation of model-generated interindividual variation. A) Empirical and generated connectomes are compared using identifiability and network measures. Identifiability is evaluated based on a similarity matrix that compares empirical and generated connectomes, and network measures can be classified into edge, node, and graph levels. B) Generated connectomes capture significant individual variability. Generated variability is measured with differential identifiability and network measures (empirical-generated Pearson correlation). Distributions show measures across 1,000 bootstraps. In all panels, *** denotes a significant (permutation test, FDR corrected *p* < 0.001) difference between model and null. Top left: differential identifiability. Top right: empirical-generated Pearson correlations in edge-level measures. Bottom left: node-level measures. Bottom right: graph-level measures.

Identifiability was used to evaluate whether an individual’s generated connectome resembles their own empirical connectome more than the ones of other individuals (Amico & Goñi, 2018). Various identifiability measures are available; we focused on the differential identifiability (Methods) and reported results for other identifiability measures in supplementary (Fig. S2). A larger differential identifiability (Amico & Goñi, 2018; Jamison et al., 2024) suggests that generated connectomes captured more empirical variation. Benchmarked to the null model, our generated connectomes achieved a significantly greater differential identifiability (model: 0.09 ± 0.01, null: 0.00 ± 0.01; permutation test, FDR-corrected *p* < 0.001; Fig 2B).

We used network measures to assess whether topological properties of an individual’s generated connectome matched those measured in their own empirical connectome. We considered twenty network measures evaluated in the literature (Rubinov & Sporns, 2010), spanning the edge, node, and graph levels (Methods). We found that network measures in our model-generated connectomes positively associated with those measures in the corresponding empirical connectomes, achieving an average correlation of Pearson *r* = 0.06 ± 0.03 across all network measures (Fig. 2B). Similar results were observed using the DK atlas (Fig. S7). In contrast, the individual variability in network measures generated by the null and MI model (Fig. S3) were smaller (null: *r* = 0.00 ± 0.00; MI: *r* = 0.01 ± 0.01).

In summary, these results demonstrate that our generative model captures significant interindividual variation in empirical connectomes. While the generated variability remains modest, it represents substantial improvement over the established state-of-the-art.

### Relative influence of personal measures to connectome generation

Next, we evaluated the relative impact of different categories of personal data to connectome generation. To this end, we iteratively suppressed categories from the generative model inputs (Methods). We defined the normalized categorical contribution (NCC, detailed in Methods), measured by the normalized reduction in generated variability when a specific category is suppressed. A larger, positive NCC suggests a category is more important to generating interindividual connectome variation.

Figure 3 shows the NCCs for each category. In this figure, Panel A) summarizes the average NCC across all identifiability and network measures. NCCs differed significantly across categories (Fig. S5), with age and sex, and body phenotypes contributing the most in the averaged NCC (41.8% and 42%, respectively), which was primarily attributed to their shared variance (Fig. S6). Panels B)-D) detail the differential identifiability, as well as example edge-level, node-level, and graph-level network measures (see all measures in Fig. S4). While the patterns of NCCs in many measures align with the averaged NCC (e.g., differential identifiability and edge-level network measures, Fig. 3B and 3C), some deviations were observed (e.g., local clustering, Fig. 3D; more in Fig. S4). Notably, lifestyle factors and cognitive measures, while being less important to the average NCCs, are crucial determinants in certain network measures (Fig. S4). In addition, a single category can enhance generated variation in one measure while constraining it in another, indicating that the model balances a trade-off between different topological aspects when generating connectomes. Observations were similar using the replication parcellation (Fig. S8).

**Figure 3.**
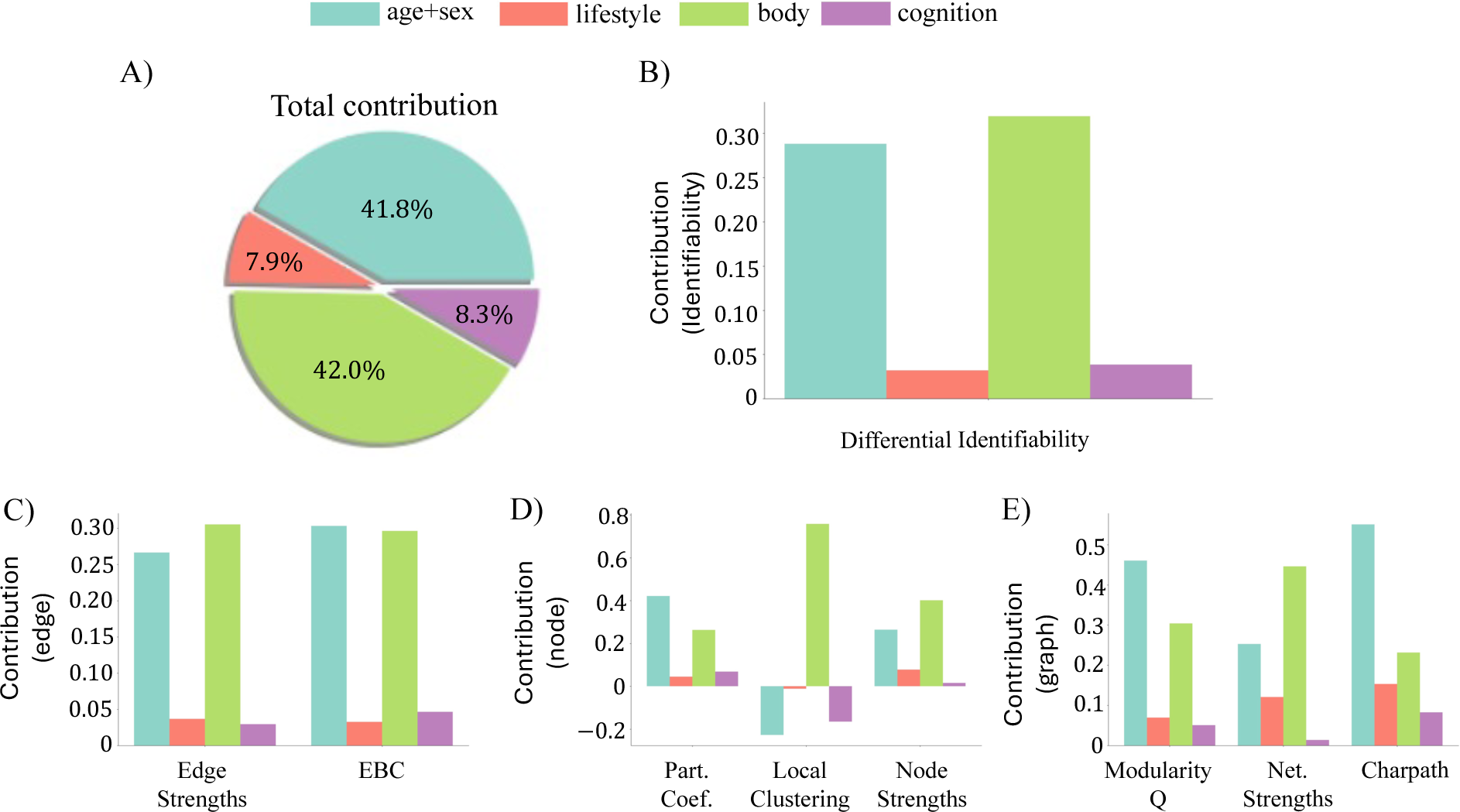
Relative importance of individual measures to connectome generation. A) Normalized categorical contribution (NCC) averaged across all network and identifiability measures. Body phenotypes and age and sex contribute more than lifestyle factors and cognitive measures. B) NCC of each personal data category to model-generated variation, measured by differential identifiability. C) Same as B) but measured by example edge-level network measures. D) Same as B) but measured by example node-level network measures. E) Same as B) but measured by example graph-level network measures.

Collectively, these results suggest that while body phenotypes and demographic characteristics are more influential predictors of connectomic variation in general, other factors such as lifestyle and cognition also contribute to shaping specific aspects of generated connectivity.

### Variability generation across functional brain subnetworks

So far, we have focused on evaluating our generative model across the entire connectome. However, the generated variability and each personal measure’s influence may vary across the brain. To investigate this, we assigned connectome regions (nodes) to 7 established functional brain subnetworks (Yeo et al., 2011).

We first assessed whether the generated variability, measured by identifiability and network measures, significantly differed between pairs of subnetworks. We found that the visual system (VIS) exhibited significantly lower (Wilcoxon test, FDR-corrected *p* < 0.05, Fig. 4A right) model-explained interindividual variation compared to the somatomotor network (SMN), default mode network (DMN), and the executive control network (ECN). Additionally, the limbic system (LIM) also observed less generated variability than the DMN. These differences were not explained by the between-subnetwork differences in interindividual variation observed in the empirical connectomes (permutation test, *p* = 10^-4^, Supplementary Fig. S9).

**Figure 4.**
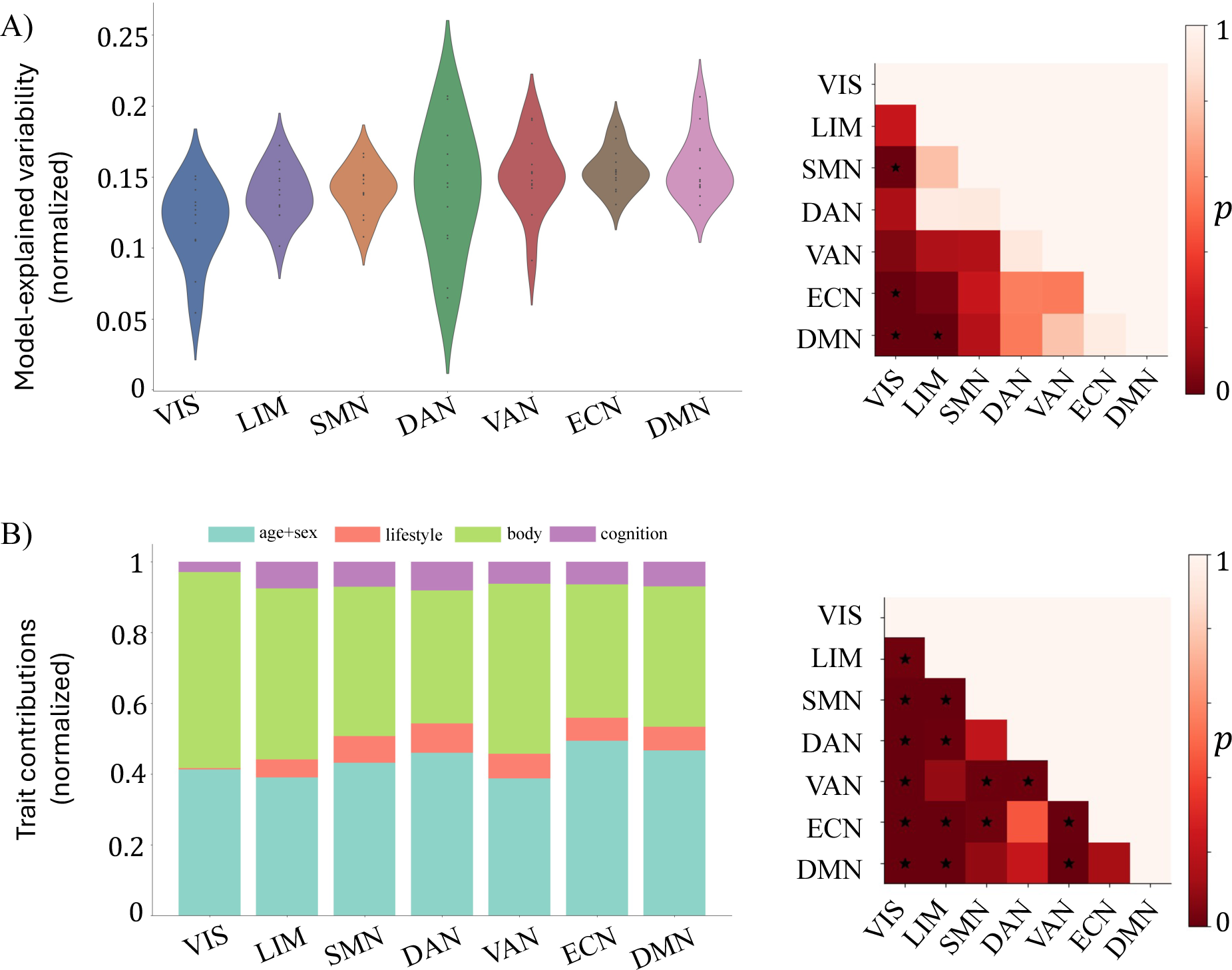
Generated variability and each category’s influence on generation heterogeneously distributed within the brain. A) Comparison of generated variability between functional subnetworks. The left figure shows violin plots of generated variability in each subnetwork (ascending order). The lower triangular matrix of the right plot displays FDR-corrected p-values for Wilcoxon tests comparing generated variation between pairs of subnetworks, with * indicating significant differences. VIS – visual system; LIM – limbic system; SMN – somatomotor network; DAN – dorsal attention network; VAN – ventral attention network; ECN – executive control network; DMN – default mode network. B) The profiles of categorical contributions in subnetworks significantly differ. The left plot visualizes the categorical contributions in each subnetwork (averaged across identifiability and network measures) in stacked bar plots, with sum of categorical contributions normalized to 1. Right figure shows the FDR-corrected p-values of paired permutation tests. * indicates significant differences between subnetworks.

Next, we evaluated whether differences in generated variability were related to differential contributions of personal data across brain subnetworks. To this end, we compared the influence of personal measures to generation between pairs of subnetworks (Methods). As shown in Fig. 4B, body phenotypes, age and sex contributed more than lifestyle and cognition across all subnetworks. However, the level of contribution from these personal factors significantly differed between 15 out of 21 pairs (permutation test, FDR-corrected *p* < 0.05, Fig. 4B right), suggesting that personal data differentially impacts connectivity generation across different brain subnetworks.

In summary, the individual variability generated by our model differs across functional brain subnetworks, which likely reflects differences in the degree to which personal profiles influence specific brain areas.

### Data augmentation using connectome generation

Finally, we aimed to demonstrate the utility of our generative model in a data augmentation task. Specifically, we used our model to generate connectomes to augment the training samples for various machine learning applications. We considered scenarios in which personal data was available for the entire cohort, but diffusion MRI data was incomplete or missing for a proportion of the cohort. Our generative model enables estimation of connectomes for individuals without diffusion MRI data.

We investigated prediction tasks commonly considered in the literature, including age prediction and sex classification (He et al., 2023; Kopetzky et al., 2024; Leming & Suckling, 2021; Sun et al., 2024). We evaluated the performance of linear and logistic regression models (L1 regularized) on these tasks. Model performance was compared between four different training data compositions: i) Empirical: all connectomes are empirically mapped, ii) Generated: all connectomes are generated from personal profiles using our model, iii) Hybrid: a balanced mixture of (i.e., equal-sized) empirical and profile-generated connectomes, and iv) Interpolation: a balanced mixture of empirical and linearly interpolated connectomes (i.e., averaging two random empirical connectomes from the training data, see Methods). The latter provides a benchmark case in which the training set is augmented by linear interpolation of existing connectomes. Models were trained with sample sizes (*N*) ranging from 8 to 512 and were evaluated with unseen test data (Fig. 5). Sample size here refers to the combined total of empirical and generated connectomes.

**Figure 5.**
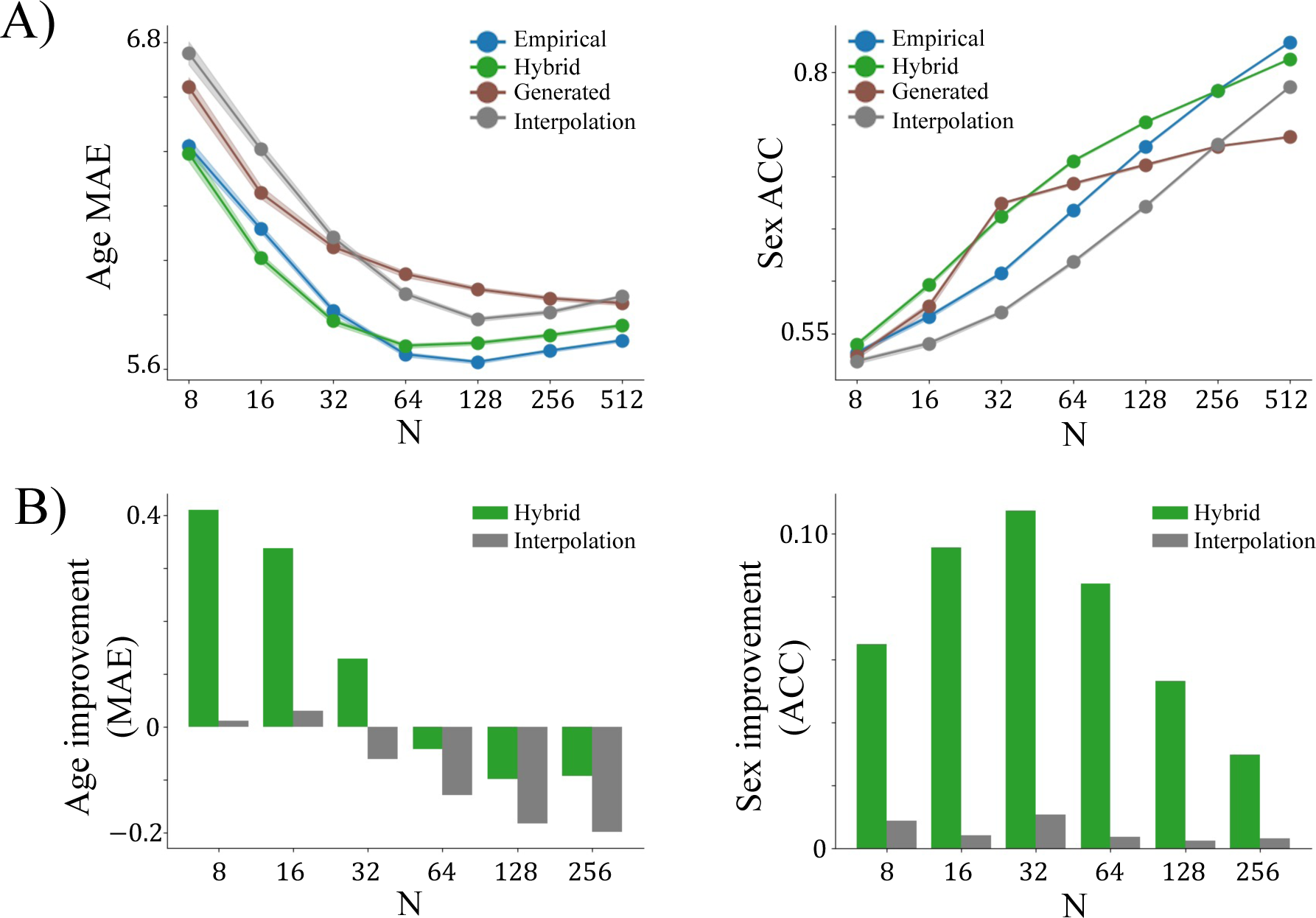
Age prediction and sex classification models trained on four types of data compositions. A) Model performance as a function of sample sizes and data compositions. Left plot displays the mean absolute error (MAE) of age prediction achieved by models. Smaller MAE indicates better performance. X-axis indicates the total number of samples in the training set. Confidence intervals (shaded areas) are derived from 1,000 bootstraps. Right figure is same as left plot but for sex classification accuracy (ACC). Larger ACC indicates better performance. B) Performance gain from data augmentation. Each bar plot displays the improved model performance achieved by an augmented training set (sample size 2*N*, including *N* empirical and *N* augmented samples), relative to the half-sized (sample size *N*) empirical samples. A positive improvement indicates a better performance (i.e., lower MAE and higher ACC) in hybrid/interpolation (2*N*) relative to empirical (*N*). Hybrid achieved stronger improvement relative to interpolation. X-axis indicates the number of empirical samples.

Increasing sample size benefited model performance in both age prediction (before plateauing near *N* = 128) and sex classification under all training data compositions. At small sample sizes (*N* < 32 for age prediction and *N* < 256 for sex classification), models trained on hybrid data performed the best (Fig. 5A). However, performance was eventually surpassed by models trained on empirical data with large sample sizes. We evaluated the performance gains of models trained on hybrid and interpolation data with a sample size of 2*N*, relative to models trained on empirical data with a sample size of *N* (Fig. 5B). This evaluated whether adding *N* profile-generated or linear augmented samples can improve the model training when researchers have access to *N* empirical samples. We found when *N* is small, including augmented data can decrease the prediction errors. Compared to interpolation, hybrid achieved a stronger improvement in model performance, with up to around 0.4-year-old smaller MAE in age prediction and more than 10% higher accuracy in sex classification (depending on sample size), relative to a conventional training sample comprising half the number of empirical connectomes (Fig. 5B). These results indicate that connectome generation can be an efficient data augmentation approach.

## Discussion

Using a person’s brain connectivity to predict their cognitive performance, behavioral measures, and other individual factors is an intensely studied problem, and numerous predictive models to achieve this goal have been established (Kawahara et al., 2017; Kopetzky et al., 2024; Lu et al., 2024; Lv et al., 2024; Seguin et al., 2020). In this work, we investigated the inverse problem of inferring a person’s brain connectivity based on their personal profile. We established a deep model that successfully generated connectomes given a person’s age, sex, and behavioral/cognitive profile. We demonstrated that our generated connectomes were able to recapitulate interindividual variation observed in connectomes empirically mapped with diffusion MRI and tractography. While the extent of individual variation generated by our model was modest, it achieves substantial improvement over the established state-of-the-art. In further analyses, we identified characteristics of an individual’s personal profile that are most important to the connectome generative process. We also demonstrated the utility of our generative model in the context of a data augmentation task, where the training sample can be augmented for individuals with missing or poorly reconstructed connectomes. Overall, we provide the first comprehensive demonstration of connectome generation based on an individual’s personal profile and provide guidance for future improvement in model design.

Mapping connectomes between modalities—structural connectivity to functional connectivity, and vice-versa—is related to the connectome generation problem considered here and is more widely explored in the literature (Chen et al., 2024; Jamison et al., 2024; Li et al., 2022; Sarwar et al., 2021; Yang et al., 2022; Zalesky et al., 2024). Generating connectomes from personal profiles is however a more challenging task than inter-modality mapping because the generation process does not have access to any brain-based information. Nevertheless, we found that the identifiability of profile-generated connectomes is comparable to state-of-the-art cross-modality connectome predictions (Jamison et al., 2024). Our generated connectomes achieved a de-meaned differential identifiability of 0.09, which compares to a range of 0.02 to 0.15 in the case of cross-modality prediction.

Connectomes are high dimensional, and capturing variation in one network property does not guarantee other aspects of connectivity and topology are similarly captured. For example, in the seminal connectome growth model that considers wiring distance and matching index (i.e., the fractional overlap in neighboring nodes) between nodes (Betzel et al., 2016), characteristic path length and modularity were significantly replicated whereas network diameter and assortativity were not. Our model captured significant interindividual variation in 19 out of 20 network measures, indicating an ability to simultaneously replicate different aspects of connectome topology.

We found that biological measures, including body phenotypes and age and sex, are typically more important contributors to connectome generation than lifestyle factors and cognition. In the literature, age and sex have strong and reproducible associations to brain connectivity and are commonly controlled as nuisance factors in neuroscience research (Geerligs et al., 2015; Ingalhalikar et al., 2014), whereas the association between connectivity and cognitive measures is relatively weaker and inconsistent (Genon et al., 2022). The observed differences across categories of individual measures provide valuable insights into which measures should be prioritized in future models when data availability is constrained by costs and missing values. Measurements with strong associations to brain connectivity, such as genetic profiles (Elliott et al., 2018; Smith et al., 2021), would enhance the model’s ability to generate greater interindividual variation in connectomes.

We observed that variation in our generated connectomes is heterogeneously distributed within the brain. Relatedly, Chen et al. (2024) reported the same observation in functional connectomes predicted from structural connectivity, where they suggest the intrinsic heterogeneity of interindividual variations in empirical connectomes may play a role. While this factor may be at play, our generated networks show a significantly different pattern of generated variability compared to empirical observations. This result suggests other factors may also contribute to the observed heterogeneity. A possible explanation unique to our generative process is that measures provided to the model may explain more variation in certain brain regions than others. For example, cognition is closely related to the association cortex (Yeo et al., 2015), and it is intuitive that cognitive measures explain more variation in the association cortex than in the visual system. Our results support this argument, with cognition showing a distinctively smaller contribution when generating connectivity for the visual system. Understanding this spatial heterogeneity in generated variability and the influence of personal measures can help improve future models. Specifically, focusing on brain regions with least model-generated variation and incorporating more relevant measures could be an efficient approach—much like how the shortest stave in a barrel determines its capacity.

The benefits of using synthetic data have been widely investigated in medical imaging research (Ibrahim et al., 2024; Khosravi et al., 2024), yet its utility in connectomic studies remains unexplored. Our work provides the first empirical evidence that generating connectomes can be an efficient data augmentation approach. Models trained on a mixture of empirical and generated data achieved promising performance for small to medium sample sizes, a phenomenon that has been observed in the generative modeling literature outside of the field of neuroscience (Bluethgen et al., 2024). This observation suggests that our generative model is a promising tool when connectivity data is limited, a common challenge in real-world studies due to collection costs, imaging artifacts, and participant dropout. In addition, in contrast to established models (Betzel et al., 2016; Liu et al., 2021), generation conditioned on personal profiles enables precise control over the generated connectivity, making the generated connectomes more relevant to the study population. This is particularly useful when certain subgroups are underrepresented in empirical data, such as rare clinical diagnoses.

Moreover, the utility of the model extends beyond data augmentation. For example, connectome research has significantly benefited from large-scale, publicly available neuroimaging datasets (Casey et al., 2018; Sudlow et al., 2015; Van Essen et al., 2013). Despite this, data privacy is a concern of data-releasing sites, and strategies that hinder data interoperability (Chakravorty et al., 2022), such as download restrictions, have been implemented. Connectome generation can promote anonymous data sharing, alleviate the conflicts between data security/privacy and accessibility in open science. However, further work is required to understand whether our generative model can facilitate data sharing under these privacy restrictions. In addition, recent research emphasizes the significance of developing tools to automate connectome quality control (Dey et al., 2022; Zalesky et al., 2024). With VAE’s capability of outlier detection (An & Cho, 2015), our model can potentially enable quality control of connectivity data with individual measures taken into consideration. These applications could be explored in future studies.

Several methodological considerations should be discussed. First, as a proof of concept, we use a simple MLP-based VAE. Recent progress in structure-function coupling studies has demonstrated that more advanced model architectures, such as graph neural networks, can better model brain connectivity (Chen et al., 2024; Li et al., 2019; Neudorf et al., 2022). Second, many studies in the literature, when modeling brain connectivity with deep neural networks, include a dispersion term in the loss function that aims to separate individuals from each other to improve identifiability (Jamison et al., 2024; Sarwar et al., 2021). In contrast, we did not include the dispersion loss in our model. This is because recent work demonstrated that dispersion-dependent improvement in identifiability is traded-off by a decrease in accuracy (Jamison et al., 2024), both of which are fundamental to connectome generation. Optimizing the model with respect to accuracy while evaluating the model with respect to interindividual variation, our results provide a conservative estimation of the empirical variation that our model can capture. Finally, while structural measures such as cortical thickness and area (Alexander-Bloch et al., 2013) are closely related to brain connectivity, we excluded them from model inputs because we aimed to model connectomes from non-brain measures. Future work could investigate to what extent structural connectivity, derived from diffusion MRI, can be informed by radiomics extracted from structural MRI.

In conclusion, our results demonstrate the feasibility of generating human connectomes from personal profiles. The generated networks replicate empirically observed individual variation in connectome network architecture, measured across numerous graph-theoretic and identifiability measures. We found that the selection of individual measures provided to the generative model is critical to the quality of generation, where biological measures are more important than behavioral/cognitive measures. Our work lays the foundation for individualized connectome generation, paving the way for future research and clinical applications.

## Methods

### Data selection and preprocessing

UKBiobank (Alfaro-Almagro et al., 2018; Sudlow et al., 2015) provides the access to individual records and MRI scans of over 40,000 participants in the middle and old lifespans. We considered individual measures of participants, categorized into age and sex, environmental and lifestyle factors, body phenotypes, and cognitive measures (Tian et al., 2023). However, not all measures were available for all individuals with structural and diffusion MRI scans. As such, we only considered participants with cognitive measures available. Environmental and lifestyle measures that were missing in more than 5% of participants were not considered. In addition, individuals with lifetime diagnosis of chronic disease (Table S1) were excluded. For participants with repeat visits, we only considered data from their initial imaging visit. These exclusion criteria result in a sample size of 8,086 participants with structural and diffusion MRIs, as well as 194 individual measures (including age, sex, 85 lifestyle factors, 78 body phenotypes, and 29 cognitive measures, see Supplementary Table S1 for measures considered) available, who were later randomly assigned into training (*n* = 6,469) and test sets (*n* = 1,617). Missing individual measures were mean-imputed and continuous variables were standardized, based on the observations in the training set.

Structural connectomes used in this study were previously mapped from structural and diffusion MRIs with probabilistic tractography (Mansour L et al., 2023). Specifically, we considered connectomes weighted by streamline counts, in the 200-node Schaefer parcellation assigned to Yeo’s 7 functional networks (Schaefer et al., 2018; Yeo et al., 2011). Connectomes in the Desikan-Killiany atlas (Desikan et al., 2006) were used to replicate the results wherever appropriate. Connectivity weights in the structural connectomes are known to span orders of magnitudes, which can lead to instable model training (Sarwar et al., 2021) and impact the computation of weighted network measures. To mitigate this, connectivity weights were log-transformed to ensure that the distribution of connectivity weights was closer to a normal distribution, a common practice that were employed in the literature (Chen et al., 2024).

### cVAE architecture and training

The cVAE model utilized in the present study included a profile projector, an encoder, and a decoder, each became a fully connected multilayer perceptron (MLP). The profile projector comprised two layers, projecting 194-dimensional individual measures *T* into 128-dimensional representation *t* with a hidden layer of 128 neurons. The encoder compressed the weights of 19,900 unique connectome edges (i.e., the upper triangular matrix) into a 128-dimensional vector representation through four feedforward layers, with a compression rate of 4 in the first three layers (layer widths [19900, 4975, 1243, 310, 128]). A single feedforward layer then summarized the combined connectome and profile representations into the mean (μ) and log variance (σ) of the latent space, from which the 128-dimensional latent representation *z* was sampled with the reparameterization trick (Kingma, 2013). Finally, the decoder reconstructed the connectome edges from the latent representation *z* and the profile representation *t* using a four-layer MLP (layer widths [256, 310, 1243, 4975, 19900]). Each layer in the encoder and decoder was followed by a layer normalization.

We treated connectivity weights as tabular data and fitted a scikit-learn StandardScaler using the training set to standardize each edge’s log-transformed weight based on its population distribution. The model was next trained on the standardized connectivity from the training set with a learning rate of 10^-4^ and batch size of 32, where an AdamW optimizer (Loshchilov, 2017) minimized the combined reconstruction and KL divergence loss, equivalent to maximizing the evidence lower bound on the log-likelihood. A dropout rate of 0.5, weight decay of 10^-4^, and early stop at 200 epochs were used during training to avoid overfitting.

### Connectome generation

Using the trained model, we generated connectomes for individuals in the test set, based on their individual profiles and a latent representation *z* randomly sampled from a standard gaussian distribution. We generated 20 networks for each individual in the test set. The amount of empirical interindividual variations generated by our model was quantified by identifiability and network measures (see below) using 1,000 bootstraps. In each bootstrap, we randomly sampled a generated network per individual (1 out of 20) to form a generated population and evaluated the extent of generated variability by comparing the generated sample with empirical connectomes.

### Null models and baselines

To establish a benchmark for the model-generated variability, we generated null networks where all information about personal data were suppressed. This was achieved by replacing all personal measures with their corresponding group average values in the training set, serving as a null model that has no access to individual data.

In addition, we generated individual connectomes using the matching index (MI) model (Betzel et al., 2016), a conventional approach governed by two parameters and considered the state-of-the-art model in connectome generation. This model simulates connectome-like network by optimizing the trade-off between cost and efficiency in brain wiring. Specifically, following Liu et al. (2023), we estimated the optimal model parameters for individuals in the training set. We then trained an L1-regularized linear regression model to predict these parameters from personal profiles and applied it to the test set. Finally, connectomes were generated for the test set using profile-predicted parameters. While the binary nature of connectomes generated from MI model limited direct comparison to our model in certain measures of interindividual variation (e.g., differential identifiability), it provided a state-of-the-art baseline for assessing model-generated variability, in other identifiability and network measures (see supplementary).

### Identifiability measures

The identifiability measures were evaluated based on the similarity matrices *S* (Fig. 2A), defined by either Pearson correlation, de-meaned correlation, or negative mean squared error, between empirical and generated connectomes. The matrix element *S_ij_* represents the similarity between the empirical connectome of subject *i* and the generated connectome of subject *j*. When constructing the similarity matrices for whole-brain connectivity (i.e., for results in Fig.2 and Fig.3), we compared connectomes using their upper triangular elements. For functional subnetworks (i.e., for results in Fig. 4), we first averaged the connectivity of all nodes within each subnetwork into a single connectivity profile vector (200 × 1 in the 200-node Schaefer parcellation). Similarity matrices were then constructed to compare these connectivity vectors between connectomes.

Different measures of identifiability that have been used in the literature, including top-1 accuracy, Hungarian accuracy, average rank percentile, and differential identifiability, were computed (Amico & Goñi, 2018; Jamison et al., 2024; Zalesky et al., 2024). The top-1 accuracy is the probability that each subject’s generated connectome is most similar to their own empirical connectome, compared to all other subjects’ empirical connectomes (Jamison et al., 2024). Note that multiple generated connectomes can be assigned to the same subject when calculate the top-1 accuracy. The Hungarian accuracy represents the probability of correct generated-empirical matching when one-to-one mapping is mandated (Zalesky et al., 2024). The average rank percentile measures the percentile rank of a generated connectome’s similarity to its own empirical counterpart, compared to other empirical connectomes (Jamison et al., 2024; Kwon et al., 2024). The differential identifiability calculates the difference in mean between the diagonal and off-diagonal elements of the similarity matrix (Amico & Goñi, 2018). For improved computational tractability, identifiability was evaluated in a subset of randomly sampled 200 subjects in each bootstrap. The expectations of top-1 accuracy, Hungarian accuracy, average rank percentile, and differential identifiability at chance level are 0.005, 0.005, 0.5, and 0, respectively. We evaluated 12 definitions of identifiability (3 similarity measures by 4 identifiability formula) and focused on the differential identifiability computed from de-meaned correlation similarity in the Results.

### Network measures

Twenty different network measures, as shown in Table 1, were evaluated for empirical and generated connectomes. The detailed interpretation of each measure can be found in Brain Connectivity Toolbox (Rubinov & Sporns, 2010).

**Table 1.** Summary of network measures. Edge level: 1-3; node level: 4-12; network level: 13-20.

| Index | Network measure | Index | Network measure |
| --- | --- | --- | --- |
| 1 | Edge weights | 11 | Nodal eccentricity |
| 2 | Edge betweenness centrality | 12 | Local Efficiency |
| 3 | Topological distance between endpoints | 13 | Global average clustering |
| 4 | Participation coefficient | 14 | Modularity Q |
| 5 | Module degree z-score | 15 | Network assortativity |
| 6 | Eigenvector centrality | 16 | Network total strengths |
| 7 | Node betweenness centrality | 17 | Characteristic path length |
| 8 | Local clustering | 18 | Global efficiency |
| 9 | Local assortativity | 19 | Network radius |
| 10 | Nodal strengths | 20 | Network diameter |

As shown in Fig. 2A, a network measure can be represented as either a matrix (edge-level), a vector (node-level), or a numeric value (graph-level). We considered the correlation in each of these network measures between the empirical and generated connectomes, where each data point in the correlation represented an individual. This correlation is referred to as the *empirical-generated correlation*, and a larger, positive empirical-generated correlation indicates a stronger model-generated variability. For node (edge) level measures, the correlation coefficient was first computed separately for each node (edge), then transformed to Fisher’s z, averaged across all nodes (edges), and finally transformed back to the correlation coefficient. This approach is known to minimize the bias when estimating the average of correlations (Corey et al., 1998). The evaluation in brain subnetworks followed the same procedure except two differences: i) the Pearson r was only averaged across nodes/edges that are involved in the subnetwork, and ii) graph-level measures were not considered because they are summary measures for the whole-brain connectome.

Special cases were carefully handled when computing the empirical-generated correlations. For example, when isolated nodes were found in empirical connectomes due to poor tractography reconstruction, its topological distance to other nodes became infinity and were considered unrealistic. These undefined network measure elements were excluded when computing the correlation. Undefined correlations were also found when a node/edge showed a constant graph-theoretical measure across all subjects, and these values were excluded when averaging across correlations. In addition, the computations of certain network measures depended on the community structure of networks, whose assignment can deviate between individual connectomes and between repetitions. To maintain the consistency of the results, we pre-computed the community structures of the group average connectivity matrix using Louvain community detection (Blondel et al., 2008; Rubinov & Sporns, 2010) with 100 independent initializations. A group consensus community assignment (Lancichinetti & Fortunato, 2012) was then derived and was used to compute the relevant network measures for individual connectomes.

### Measuring categorical contributions of personal data

To resolve the influence of individual profile categories to connectome generation, we further generated networks where individual measures were iteratively suppressed by category. We defined the normalized categorical contribution (NCC) to quantify the relative influence of category *i* to variability generation, measured by each network or identifiability measure, as

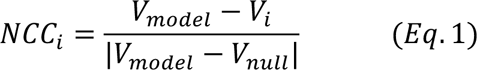

where *V_model_*, *V_i_*, and *V_null_* represent the generated variability in model connectomes (i.e., no individual measures suppressed), category *i* suppressed connectomes, and null connectomes (i.e., all personal profiles suppressed), respectively. In brief, *NCC_i_* quantifies the drop in generated variability when category *i* is suppressed, normalized by the difference between the profile-complete model and null. The absolute value operator is included to ensure that a positive *NCC_i_* indicates measure category *i* improves generated variability, even if *V_model_* < *V_null_*.

### Generated variability and categorical contributions across functional brain subnetworks

To evaluate whether the generated variation varied across the brain, we quantified the generated variability in each of the seven functional brain subnetworks defined by Yeo et al. (2011), as described in *Methods –identifiability/network measures*. Generated variability was assessed using 3 identifiability measures and 10 network measures. Identifiability measures are generally similar to each other; thus, we only considered differential identifiability because they are continuous (three definitions of similarity). Two node-level measures, local clustering and local efficiency, were excluded because their empirical-generated correlations were weak (*r* < 0.02). Such low correlations can result in high coefficient of variation (i.e., the ratio of mean Pearson r to the standard deviation), leading to unreliable estimation of generated variability in subnetworks. A Wilcoxon test was used to assess whether the generated variability significantly differed between pairs of subnetworks.

We quantified the relative influence of personal data categories across subnetworks to determine whether they differed. A schematic illustration of this process can be found in Fig. S10. A subnetwork was represented by 13 vectors, each corresponding to an identifiability and network measure considered. Vector elements were NCCs that quantified the categorical contributions to generation, as described in *Methods – Measuring categorical contributions of personal data*. We compared the dissimilarity of vectors within and between subnetworks. Specifically, for each pair of subnetworks, the average Euclidean distance between vectors defined *D_within_* (i.e., dissimilarity of vectors within the subnetwork) and **D*_between_*, (i.e., dissimilarity of vectors from different networks). The value of *D_between_*, − *D_within_*, was benchmarked to a null distribution derived from paired permutation tests. Permutations were restricted such that vector shuffling only occurred within the same network or identifiability measure, resulting in 2^12^ = 4096 possible combinations. The p-value was calculated as the proportion of permuted combinations where the between-minus-within difference exceeded the empirically observed results. A significant result indicated that the contribution of personal data to connectome generation differs more between than within subnetworks, supporting the presence of distinct contribution profiles across the brain.

### Data augmentation

To investigate the application of data augmentation through connectome generation, we implemented linear and logistic regression models with L1 regularization using scikit-learn, to predict individual age and sex from connectivity. At a sample size 2*N*, we randomly sampled 2*N* subjects from the 1,617 individuals in the test set. The empirical (generated) training data was composed of the empirical (profile-generated) connectomes of the 2*N* subjects. The augmented (hybrid/interpolation) training data was composed of *N* empirical and *N* augmented connectomes. In hybrid, augmented connectomes were generated from personal data. In interpolation, augmented connectomes were generated by averaging the connectomes of two randomly sampled subjects with the same sex (from the *N* empirical subjects), and the ground truth label in age was set to the averaged age. Age prediction MAE and sex classification ACC were evaluated with the empirical data of unselected subjects in the 1,617 test set individuals.

## Code and data availability

The personal records, neuroimages, and structural connectomes used in the present study can be accessed via the UK Biobank Access Management System (https://ams.ukbiobank.ac.uk/ams/). Connectomes were previously mapped by S.M.L and returned to UK Biobank (https://github.com/sina-mansour/UKB-connectomics). The code for conducting analyses is available on GitHub (https://github.com/yuanzhel94/generate_personalized_connectomes).

## Acknowledgements

This research is conducted using data from the UK Biobank. The computation is supported by the University of Melbourne’s Research Computing Services and the Petascale Campus Initiative.

## References

Akarca, D., Vértes, P. E., Bullmore, E. T., & Astle, D. E. (2021). A generative network model of neurodevelopmental diversity in structural brain organization. Nature communications, 12(1), 1–18.

Alexander-Bloch, A., Giedd, J. N., & Bullmore, E. (2013). Imaging structural co-variance between human brain regions. Nature reviews neuroscience, 14(5), 322–336.

Alfaro-Almagro, F., Jenkinson, M., Bangerter, N. K., Andersson, J. L., Griffanti, L., Douaud, G., Sotiropoulos, S. N., Jbabdi, S., Hernandez-Fernandez, M., & Vallee, E. (2018). Image processing and Quality Control for the first 10,000 brain imaging datasets from UK Biobank. Neuroimage, 166, 400–424.

Amico, E., & Goñi, J. (2018). The quest for identifiability in human functional connectomes. Scientific reports, 8(1), 8254.

An, J., & Cho, S. (2015). Variational autoencoder based anomaly detection using reconstruction probability. Special lecture on IE, 2(1), 1–18.

Betzel, R. F., Avena-Koenigsberger, A., Goñi, J., He, Y., De Reus, M. A., Griffa, A., Vértes, P. E., Mišic, B., Thiran, J.-P., & Hagmann, P. (2016). Generative models of the human connectome. Neuroimage, 124, 1054–1064.

Blondel, V. D., Guillaume, J.-L., Lambiotte, R., & Lefebvre, E. (2008). Fast unfolding of communities in large networks. Journal of Statistical Mechanics: Theory and Experiment, 2008(10), P10008.

Bluethgen, C., Chambon, P., Delbrouck, J.-B., van der Sluijs, R., Połacin, M., Zambrano Chaves, J. M., Abraham, T. M., Purohit, S., Langlotz, C. P., & Chaudhari, A. S. (2024). A vision–language foundation model for the generation of realistic chest x-ray images. Nature Biomedical Engineering, 1-13.

Bobyleva, A., Gorsky, A., Nechaev, S., Valba, O., & Pospelov, N. (2024). Metric structural human connectomes: localization and multifractality of eigenmodes. arXiv preprint arXiv:2405.17349.

Bullmore, E., & Sporns, O. (2012). The economy of brain network organization. Nature reviews neuroscience, 13(5), 336–349.

Carozza, S., Holmes, J., Vértes, P. E., Bullmore, E., Arefin, T., Pugliese, A., Zhang, J., Kaffman, A., Akarca, D., & Astle, D. (2022). Early adversity changes the economic conditions of structural brain network organisation. bioRxiv.

Casey, B. J., Cannonier, T., Conley, M. I., Cohen, A. O., Barch, D. M., Heitzeg, M. M., Soules, M. E., Teslovich, T., Dellarco, D. V., & Garavan, H. (2018). The adolescent brain cognitive development (ABCD) study: imaging acquisition across 21 sites. Developmental cognitive neuroscience, 32, 43–54.

Chakravorty, N., Sharma, C. S., Molla, K. A., & Pattanaik, J. K. (2022). Open science: Challenges, possible solutions and the way forward. Proceedings of the Indian National Science Academy, 88(3), 456–471.

Chen, P., Yang, H., Zheng, X., Jia, H., Hao, J., Xu, X., Li, C., He, X., Chen, R., & Okubo, T. S. (2024). Group-common and individual-specific effects of structure–function coupling in human brain networks with graph neural networks. Imaging Neuroscience, 2, 1–21.

Corey, D. M., Dunlap, W. P., & Burke, M. J. (1998). Averaging correlations: Expected values and bias in combined Pearson rs and Fisher’s z transformations. The Journal of general psychology, 125(3), 245–261.

Desikan, R. S., Ségonne, F., Fischl, B., Quinn, B. T., Dickerson, B. C., Blacker, D., Buckner, R. L., Dale, A. M., Maguire, R. P., & Hyman, B. T. (2006). An automated labeling system for subdividing the human cerebral cortex on MRI scans into gyral based regions of interest. Neuroimage, 31(3), 968–980.

Dey, P., Zhang, Z., & Dunson, D. B. (2022). Outlier detection for multi-network data. Bioinformatics, 38(16), 4011–4018.

Elliott, L. T., Sharp, K., Alfaro-Almagro, F., Shi, S., Miller, K. L., Douaud, G., Marchini, J., & Smith, S. M. (2018). Genome-wide association studies of brain imaging phenotypes in UK Biobank. Nature, 562(7726), 210–216.

Feng, G., Chen, R., Zhao, R., Li, Y., Ma, L., Wang, Y., Men, W., Gao, J., Tan, S., & Cheng, J. (2023). Longitudinal development of the human white matter structural connectome and its association with brain transcriptomic and cellular architecture. Communications Biology, 6(1), 1257.

Geerligs, L., Renken, R. J., Saliasi, E., Maurits, N. M., & Lorist, M. M. (2015). A brain-wide study of age-related changes in functional connectivity. Cerebral cortex, 25(7), 1987–1999.

Genon, S., EickhoZ, S. B., & Kharabian, S. (2022). Linking interindividual variability in brain structure to behaviour. Nature reviews neuroscience, 23(5), 307–318.

Griffa, A., Baumann, P. S., Thiran, J.-P., & Hagmann, P. (2013). Structural connectomics in brain diseases. Neuroimage, 80, 515–526.

Hagmann, P. (2005). From diffusion MRI to brain connectomics.

He, Y., Chan, Y. H., & Rajapakse, J. C. (2023). Predicting gender from structural and functional connectomes via brain and population graph neural networks. bioRxiv, 2023.2011. 2001.565175.

Ibrahim, M., Khalil, Y. A., Amirrajab, S., Sun, C., Breeuwer, M., Pluim, J., Elen, B., Ertaylan, G., & Dumontier, M. (2024). Generative AI for Synthetic Data Across Multiple Medical Modalities: A Systematic Review of Recent Developments and Challenges. arXiv preprint arXiv:2407.00116.

Ingalhalikar, M., Smith, A., Parker, D., Satterthwaite, T. D., Elliott, M. A., Ruparel, K., Hakonarson, H., Gur, R. E., Gur, R. C., & Verma, R. (2014). Sex differences in the structural connectome of the human brain. Proceedings of the national academy of sciences, 111(2), 823–828.

Jamison, K. W., Gu, Z., Wang, Q., Sabuncu, M. R., & Kuceyeski, A. (2024). Release the Krakencoder: A unified brain connectome translation and fusion tool. bioRxiv.

Kawahara, J., Brown, C. J., Miller, S. P., Booth, B. G., Chau, V., Grunau, R. E., Zwicker, J. G., & Hamarneh, G. (2017). BrainNetCNN: Convolutional neural networks for brain networks; towards predicting neurodevelopment. Neuroimage, 146, 1038–1049.

Khosravi, B., Li, F., Dapamede, T., Rouzrokh, P., Gamble, C. U., Trivedi, H. M., Wyles, C. C., Sellergren, A. B., Purkayastha, S., & Erickson, B. J. (2024). Synthetically enhanced: unveiling synthetic data’s potential in medical imaging research. EBioMedicine, 104.

Kingma, D. P. (2013). Auto-encoding variational bayes. arXiv preprint arXiv:1312.6114.

Kopetzky, S. J., Li, Y., Kaiser, M., Butz-Ostendorf, M., & Initiative, A. s. D. N. (2024). Predictability of intelligence and age from structural connectomes. PloS one, 19(4), e0301599.

Kwon, J., Seo, J., Wang, H., Moon, T., Yoo, S., & Cha, J. (2024). Predicting task-related brain activity from resting-state brain dynamics with fMRI Transformer. bioRxiv, 2024.2005. 2029.596544.

Lancichinetti, A., & Fortunato, S. (2012). Consensus clustering in complex networks. Scientific reports, 2(1), 336.

Leming, M., & Suckling, J. (2021). Deep learning for sex classification in resting-state and task functional brain networks from the UK Biobank. Neuroimage, 241, 118409.

Li, Y., Mateos, G., & Zhang, Z. (2022). Learning to model the relationship between brain structural and functional connectomes. IEEE Transactions on Signal and Information Processing over Networks, 8, 830–843.

Li, Y., Shafipour, R., Mateos, G., & Zhang, Z. (2019). Mapping brain structural connectivities to functional networks via graph encoder-decoder with interpretable latent embeddings. 2019 IEEE Global Conference on Signal and Information Processing (GlobalSIP).

Liu, M., Zhang, Z., & Dunson, D. B. (2021). Graph auto-encoding brain networks with applications to analyzing large-scale brain imaging datasets. Neuroimage, 245, 118750.

Liu, Y., Seguin, C., Betzel, R. F., Han, D., Akarca, D., Di Biase, M. A., & Zalesky, A. (2024). A generative model of the connectome with dynamic axon growth. Network Neuroscience, 1-47.

Liu, Y., Seguin, C., Mansour, S., Oldham, S., Betzel, R., Di Biase, M. A., & Zalesky, A. (2023). Parameter estimation for connectome generative models: Accuracy, reliability, and a fast parameter fitting method. Neuroimage, 270, 119962.

Loshchilov, I. (2017). Decoupled weight decay regularization. arXiv preprint arXiv:1711.05101.

Lu, J., Yan, T., Yang, L., Zhang, X., Li, J., Li, D., Xiang, J., & Wang, B. (2024). Brain fingerprinting and cognitive behavior predicting using functional connectome of high inter-subject variability. Neuroimage, 295, 120651.

Lv, Q., Wang, X., Wang, X., Ge, S., & Lin, P. (2024). Connectome-based prediction modeling of cognitive control using functional and structural connectivity. Brain and Cognition, 181, 106221.

Mansour L, S., Di Biase, M. A., Smith, R. E., Zalesky, A., & Seguin, C. (2023). Connectomes for 40,000 UK Biobank participants: a multi-modal, multi-scale brain network resource. Neuroimage, 283, 120407.

Mansour L, S., Tian, Y., Yeo, B. T., Cropley, V., & Zalesky, A. (2021). High-resolution connectomic fingerprints: Mapping neural identity and behavior. Neuroimage, 229, 117695.

Neudorf, J., Kress, S., & Borowsky, R. (2022). Structure can predict function in the human brain: a graph neural network deep learning model of functional connectivity and centrality based on structural connectivity. Brain Structure and Function, 227(1), 331–343.

Oldham, S., Fornito, A., & Ball, G. (2024). Coming up short: generative network models fail to accurately capture long-range connectivity. bioRxiv, 2024.2011. 2018.624192.

Oldham, S., Fulcher, B. D., Aquino, K. M., Arnatkeviciute, A. M., Paquola, C., Shishegar, R., & Fornito, A. (2021). Modeling spatial, developmental, physiological, and topological constraints on human brain connectivity. bioRxiv.

Puxeddu, M. G., Faskowitz, J., Betzel, R. F., Petti, M., Astolfi, L., & Sporns, O. (2020). The modular organization of brain cortical connectivity across the human lifespan. Neuroimage, 218, 116974.

Rubinov, M., & Sporns, O. (2010). Complex network measures of brain connectivity: uses and interpretations. Neuroimage, 52(3), 1059–1069.

Sarwar, T., Tian, Y., Yeo, B. T., Ramamohanarao, K., & Zalesky, A. (2021). Structure-function coupling in the human connectome: A machine learning approach. Neuroimage, 226, 117609.

Schaefer, A., Kong, R., Gordon, E. M., Laumann, T. O., Zuo, X.-N., Holmes, A. J., EickhoZ, S. B., & Yeo, B. T. (2018). Local-global parcellation of the human cerebral cortex from intrinsic functional connectivity MRI. Cerebral cortex, 28(9), 3095–3114.

Seguin, C., Tian, Y., & Zalesky, A. (2020). Network communication models improve the behavioral and functional predictive utility of the human structural connectome. Network Neuroscience, 4(4), 980–1006.

Simpson, S. L., Hayasaka, S., & Laurienti, P. J. (2011). Exponential random graph modeling for complex brain networks. PloS one, 6(5), e20039.

Smith, S. M., Douaud, G., Chen, W., Hanayik, T., Alfaro-Almagro, F., Sharp, K., & Elliott, L. T. (2021). An expanded set of genome-wide association studies of brain imaging phenotypes in UK Biobank. Nature neuroscience, 24(5), 737–745.

Smolders, L., De Baene, W., Rutten, G.-J., van der Hofstad, R., & Florack, L. (2024). Can structure predict function at individual level in the human connectome? Brain Structure and Function, 1-15.

Song, H. F., Kennedy, H., & Wang, X.-J. (2014). Spatial embedding of structural similarity in the cerebral cortex. Proceedings of the national academy of sciences, 111(46), 16580–16585.

Sporns, O., Tononi, G., & Kötter, R. (2005). The human connectome: a structural description of the human brain. PLoS computational biology, 1(4), e42.

Sudlow, C., Gallacher, J., Allen, N., Beral, V., Burton, P., Danesh, J., Downey, P., Elliott, P., Green, J., & Landray, M. (2015). UK biobank: an open access resource for identifying the causes of a wide range of complex diseases of middle and old age. PLoS medicine, 12(3), e1001779.

Sun, H., Mehta, S., Khaitova, M., Cheng, B., Hao, X., Spann, M., & Scheinost, D. (2024). Brain age prediction and deviations from normative trajectories in the neonatal connectome. Nature communications, 15(1), 10251.

Tian, Y. E., Cropley, V., Maier, A. B., Lautenschlager, N. T., Breakspear, M., & Zalesky, A. (2023). Heterogeneous aging across multiple organ systems and prediction of chronic disease and mortality. Nature medicine, 29(5), 1221–1231.

Tunç, B., Solmaz, B., Parker, D., Satterthwaite, T. D., Elliott, M. A., Calkins, M. E., Ruparel, K., Gur, R. E., Gur, R. C., & Verma, R. (2016). Establishing a link between sex-related differences in the structural connectome and behaviour. Philosophical Transactions of the Royal Society B: Biological Sciences, 371(1688), 20150111.

van den Heuvel, M. P., & Sporns, O. (2019). A cross-disorder connectome landscape of brain dysconnectivity. Nature reviews neuroscience, 20(7), 435–446.

Van Essen, D. C., Smith, S. M., Barch, D. M., Behrens, T. E., Yacoub, E., Ugurbil, K., & Consortium, W.-M. H. (2013). The WU-Minn human connectome project: an overview. Neuroimage, 80, 62–79.

Vértes, P. E., Alexander-Bloch, A. F., Gogtay, N., Giedd, J. N., Rapoport, J. L., & Bullmore, E. T. (2012). Simple models of human brain functional networks. Proceedings of the national academy of sciences, 109(15), 5868–5873.

Yang, H., Winter, S., Zhang, Z., & Dunson, D. (2022). Interpretable AI for relating brain structural and functional connectomes. arXiv preprint arXiv:2210.05672.

Yeo, B. T., Krienen, F. M., EickhoZ, S. B., Yaakub, S. N., Fox, P. T., Buckner, R. L., Asplund, C. L., & Chee, M. W. (2015). Functional specialization and flexibility in human association cortex. Cerebral cortex, 25(10), 3654–3672.

Yeo, B. T., Krienen, F. M., Sepulcre, J., Sabuncu, M. R., Lashkari, D., Hollinshead, M., Roffman, J. L., Smoller, J. W., Zöllei, L., & Polimeni, J. R. (2011). The organization of the human cerebral cortex estimated by intrinsic functional connectivity. Journal of neurophysiology.

Yeung, H. W., Stolicyn, A., Buchanan, C. R., Tucker-Drob, E. M., Bastin, M. E., Luz, S., McIntosh, A. M., Whalley, H. C., Cox, S. R., & Smith, K. (2023). Predicting sex, age, general cognition and mental health with machine learning on brain structural connectomes. Human brain mapping, 44(5), 1913–1933.

Yeung, H. W., Stolicyn, A., Shen, X., Adams, M. J., Romaniuk, L., Thng, G., Buchanan, C. R., Tucker-Drob, E. M., Bastin, M. E., & McIntosh, A. M. (2024). Classification accuracy of structural and functional connectomes across different depressive phenotypes. Imaging Neuroscience, 2, 1–24.

Zalesky, A., Sarwar, T., Tian, Y., Liu, Y., Yeo, B. T., & Ramamohanarao, K. (2024). Predicting an individual’s functional connectivity from their structural connectome: Evaluation of evidence, recommendations, and future prospects. Network Neuroscience, 1-18.

Zhang, X., Braun, U., Harneit, A., Zang, Z., Geiger, L. S., Betzel, R. F., Chen, J., Schweiger, J. I., Schwarz, K., & Reinwald, J. R. (2021). Generative network models of altered structural brain connectivity in schizophrenia. Neuroimage, 225, 117510.

Zhang, Y., Liu, M., Zhang, Z., & Dunson, D. (2024). Motion-invariant variational autoencoding of brain structural connectomes. Imaging Neuroscience, 2, 1–27.

