## supplementary_information for "Deep generation of personalized connectomes based on individual attributes"

### Deep generation of personalized connectomes based on individual attributes
#### Supplementary Information

Yuanzhe Liu [a,b], Caio Seguin [a], Sina Mansour L. [a,c],
Ye Ella Tian [a], Maria A. Di Biase [a,d,e], Andrew Zalesky [a,b]

a. Systems Lab, Department of Psychiatry, The University of Melbourne, Melbourne, VIC, Australia
b. Department of Biomedical Engineering, Faculty of Engineering & Information Technology, The University of Melbourne, Melbourne, VIC, Australia
c. Center for Sleep & Cognition & Center for Translational Magnetic Resonance Research, Yong Loo Lin School of Medicine, National University of Singapore, Singapore
d. Stem Cell Disease Modelling Lab, Department of Anatomy and Physiology, The University of Melbourne, Melbourne, VIC, Australia
e. Department of Psychiatry, Brigham and Women's Hospital, Harvard Medical School, Boston, MA, USA

##### Correspondence
Yuanzhe Liu
Andrew Zalesky

##### *Generated connectomes resemble empirically observed group differences*

Generating connectomes for individuals in the test set, we observed that between-group differences evident in the empirical data were reflected in the generated samples. In Fig. 1D, we displayed the empirical and generated sex differences for test set subjects. Here we showed that empirical connectivity differences associated with age were also generated by the model. Specifically, we selected 100 youngest and 100 oldest subjects from the test set to form two subject groups and generated a connectome for each selected individual. The group average connectomes were computed for each group, and we defined the between-group difference as the averaged connectome of the younger population minus the averaged connectome of the older population. The empirical and generated between-group differences as well as their edgewise associations were shown in Fig S1.

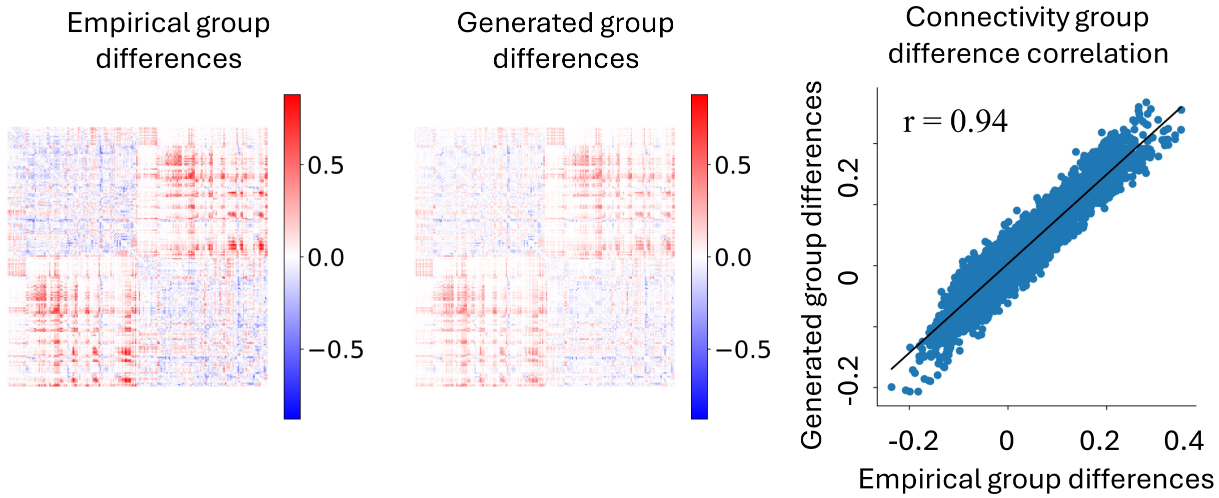


Fig S1. Generated connectomes resemble the age effects on connectivity differences between groups. Left shows the empirically observed young-old population between-group differences. Center visualizes the between-group differences in generated samples. Right compares the empirical and generated between-group differences for all edges. Each dot represents a unique edge.

##### *Network measures and abbreviations*

In this study, we comprehensively evaluated 20 network measures that have been investigated in the connectomic literature. While we focused on example measures in the main, we report results for all network measures in this supplementary document. For better clarity, we define abbreviations for network measures in supplementary table 2.

*Supplementary table 2. Network measures and their abbreviations.*

| Abbr. | Metric name | Abbr. | Metric name |
|---|---|---|---|
| $S_E$ | Edge weights | $ne$ | Nodal eccentricity |
| $ebc$ | Edge betweenness centrality | $le$ | Local Efficiency |
| $td$ | Topological distance between endpoints | $gc$ | Global average clustering |
| $pc$ | Participation coefficient | $Q$ | Modularity Q |
| $mdz$ | Module degree z-score | $r$ | Network assortativity |
| $ec$ | Eigenvector centrality | $S_G$ | Network total strengths |
| $nbc$ | Node betweenness centrality | $cpl$ | Characteristic path length |
| $lc$ | Local clustering | $ge$ | Global efficiency |
| $la$ | Local assortativity | $R$ | Network radius |
| $S_N$ | Nodal strengths | $D$ | Network diameter |

***Model outperforms null in generating individual variability***

In this work, we started by investigating if generated connectomes can preserve interindividual variations that are observed in empirical connectomes. In Figure 2 of the main, we visualized model- and null-generated variability, measured by example identifiability and network measures. Here in Figure S2, we showed the results for all measures. Note that the identifiability of model-generated connectomes is comparable to state-of-the-art cross-modality connectome predictions (Jamison et al., 2024), achieving a marginally smaller correlation average rank percentile (Fig S2D, 0.66 in our model versus 0.67 to 0.99 in cross-modality prediction depending on “connectivity flavors”, i.e., the choice of parcellations and construction pipelines) and a similar de-meaned correlation differential identifiability (Fig. S2C, center, 0.09 in profile-generation versus 0.02 to 0.15 in cross-modality prediction). These results are significant because generating connectomes from individual profiles is considered a more challenging problem than cross-modality prediction.

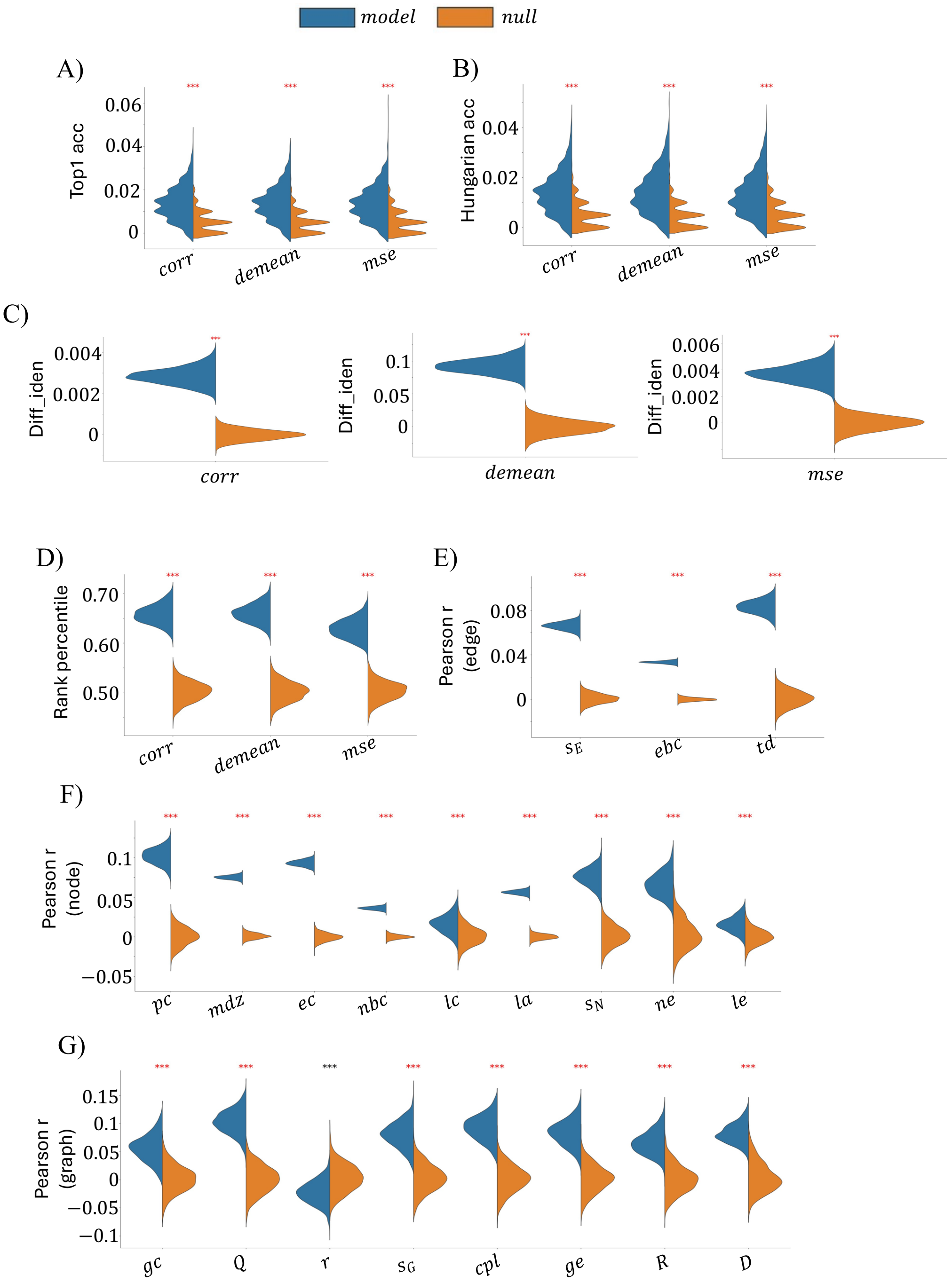


*Figure S2. Model outperforms null in generating individual variations. A)-D) Identifiability measured in top1 accuracy, Hungarian accuracy, differential identifiability, and average rank percentile, respectively, where similarity matrices are constructed using Pearson correlation, de-meaned correlation, and negative MSE, respectively. E) Empirical-generated correlation Pearson r in edge-level measures. Network measure abbreviations on the x-axis can be found in Table. S1. F) Same as E) but in node-level measures. G) Same as E) but in graph-level measures. *** mark in red color indicates significantly larger Pearson r than null, whereas *** mark in black color suggests significantly larger Pearson r in null than model (p<0.001). In network assortativity (r in Panel G), null explains greater individual variability than model. In all other measures, model explains greater individual variability than null.*

***Individual variability generated by matching index model***

We also employed the matching index (MI) model to generate individual connectomes based on their personal profile, serving as a benchmark to the deep generative model. These models are governed by two parameters, generating networks by balancing the cost-efficiency tradeoff of the human brain wiring (Betzel et al., 2016). As the classical MI model generates binary connectomes and is hard to scale with high-resolution parcellations and dense connectomes, we used connectomes in the DK atlas (68 nodes) and binarized all connectomes to a network density of 10%, a common practice in the literature (Akarca et al., 2021; Oldham et al., 2021; Zhang et al., 2021).

We first fitted the optimized model parameters for each individual using the method described in Liu et al. (2023). Next, we trained a L1 regularized linear model to predict the two optimized model parameters for individuals from their personal profiles using the training set. Finally, we predicted the model parameters for individual in the test set and generated individual connectomes with predicted parameters. We evaluated the generated individual variability using identifiability and network measures. Note that a few modifications, relative to the procedures used to evaluate weighted conenctomes, were implemented due to the binary nature of MI generated connectomes. First, for all identifiability measures, the similarity matrix between empirical and generated networks were evaluated using the Jaccard Index, which measures the percentage of overlap between two networks. Jaccard Index was also used to evaluate the existence of individual edges, analogous to the edge strengths measure (edge-level) in weighted connectomes. Finally, network measures that cannot be generalized to binary networks, including local assortativity and network total strengths, were removed from the analysis. Results were benchmarked to a binary MI null in which all individual connectomes were generated with the group average MI model parameters of the training set. Model-generated variability, as measured by identifiability and network measures, can be found in Fig. S3.

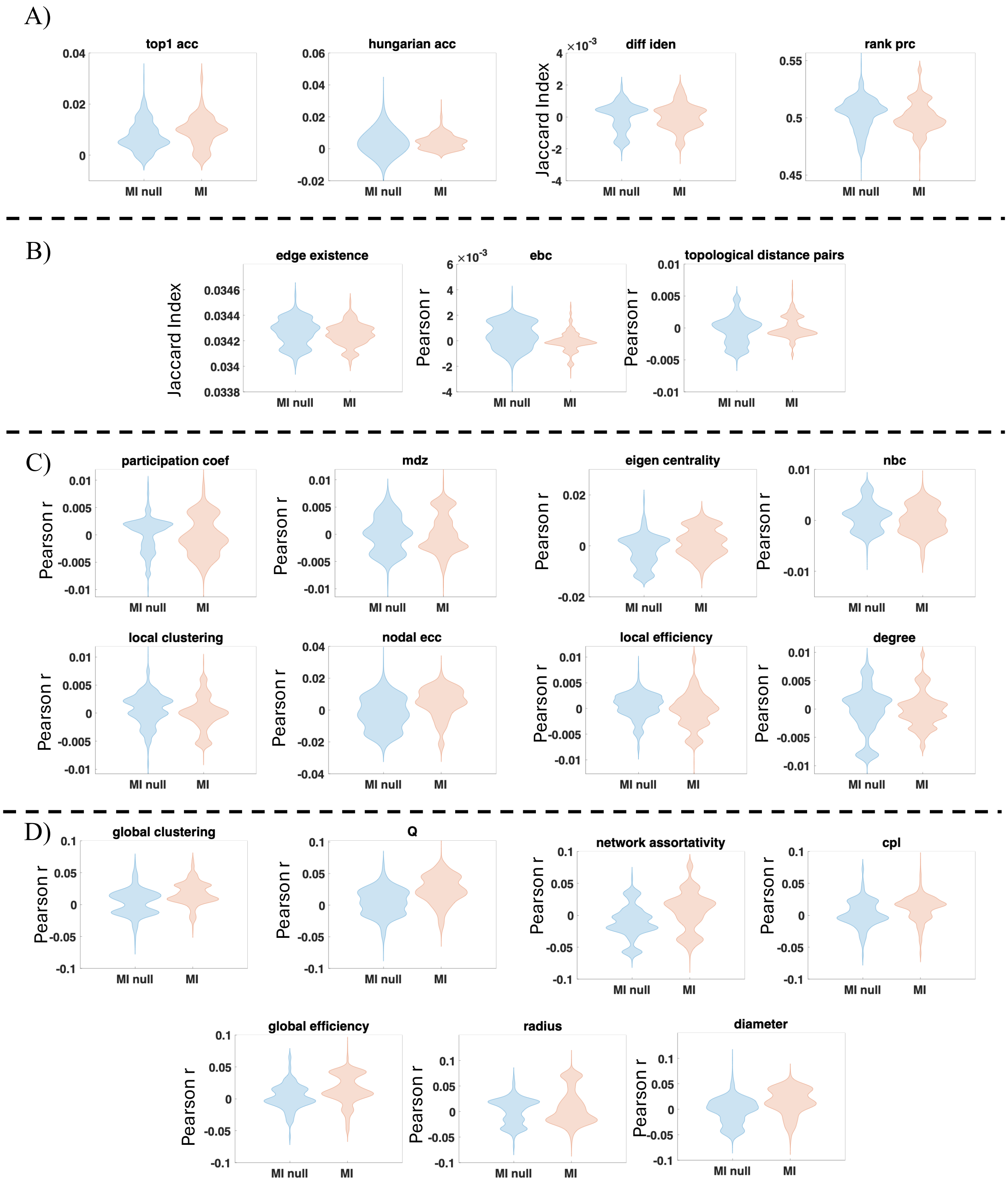


*Figure S3. MI model generate weak individual variability. A) Identifiability measures of connectomes generated by MI model and MI null. While the differential identifiability of MI model cannot be directly compared to the values of the deep model due to the binary nature of generated connectomes, other identifiability measures are all smaller than the deep model (e.g., average rank percentile of MI:* $0.50 \pm 0.01$*).B) edge-level network measures. C) node-level network measures. D) Graph-level network measures.*

#### ***Relative categorical contribution of personal data to generating variability***

In the main text, Figure 3 shows the normalized categorical contributions (NCC) for example network and identifiability measures. Here in Figure S4, we disclose the NCC for all measures. Note that while NCC in most measures align with average pattern (Fig. 3), deviation from the average can also be found.

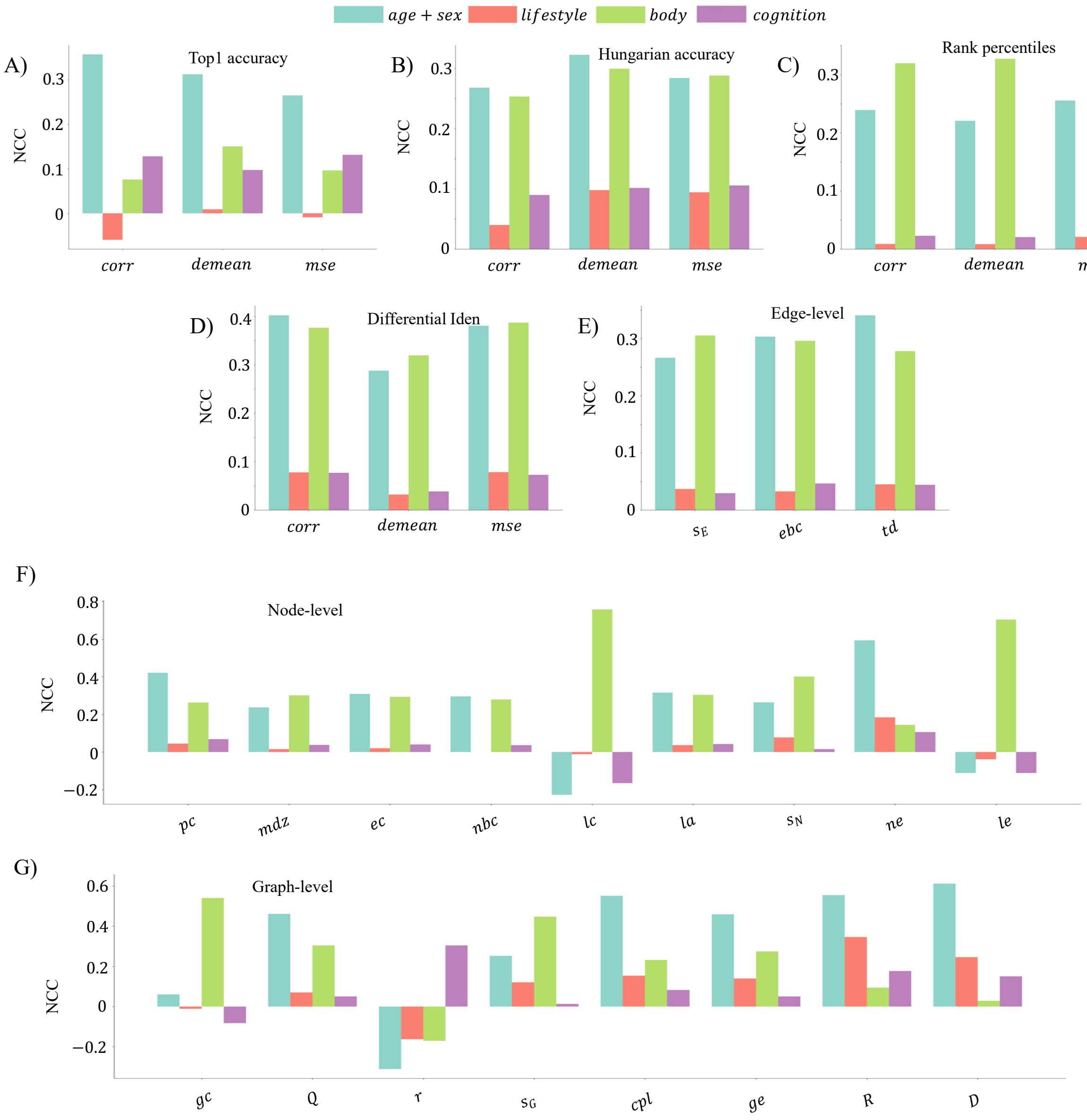


*Figure S4. NCC for all measures. A)-D) NCC in identifiability measures, including top-1 accuracy (A), Hungarian accuracy (B), differential identifiability (C), and average rank percentiles (D). E) NCC in edge-level measures. See Table. S1 for measure abbreviations on the x-axis. F) Same as E) but for node-level measures. G) Same as E) but for graph-level measures.*

##### ***Categorical contributions to model performance significantly differ***

In Figure 3 and Supplementary Figure S4, we showed the results of contributions of each personal data category, measured by 12 identifiability measures (3 similarity definition x 4 identifiability variants), 3 edge-level, 9 node-level, and 8 graph-level network measures. The significance of the contribution difference between categories were evaluated by a permutation test that compare variability generated by model (i.e., profile-complete), a single category suppressed model, and null (i.e., all categories suppressed). We found that all categories contribute significantly to model-generated variation, and the strength of contribution tend to significantly differ between categories (Figure S5).

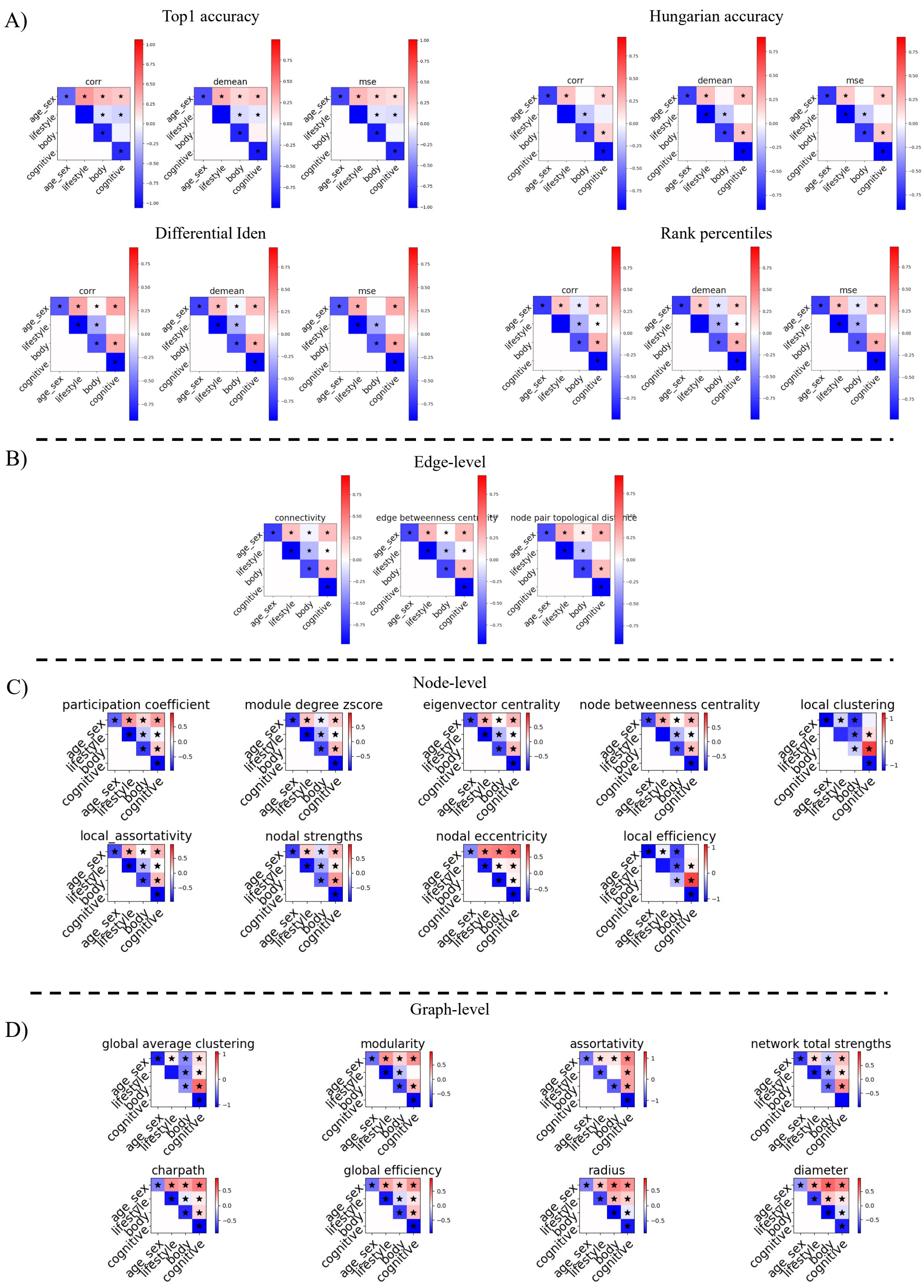


*Figure S5. Significance of categorical contributions. The diagonal elements indicate significance of categorical contribution by comparing the category-suppressed model against the category-complete model. The upper triangular elements of the matrices compare contributions between categories on the rows and columns. Blue color on diagonal suggests removing the category reduced model-generated variation; blue (red) color off diagonal suggest the category on the row contributes less (more) to model-generated variation than the category on the column. * sign denotes significant difference. A) describes the results from identifiability measured in top1 accuracy, Hungarian accuracy, differential identifiability, and average rank percentiles. B) – D) visualizes the results from edge-level, node-level, and network-level graph theory measures, respectively.*

##### ***Shared variance between body phenotypes and age and sex***

We found that body phenotypes, and age and sex exert the strongest influence on generating interindividual variation. However, these measures are interrelated, and it is unknown to which extent their shared and unique variance contribute to the generative process. To investigate this, we regressed out age and sex from other individual measures and train an independent cVAE for connectome generation, using the same network architecture and hyperparameters. Notably, this approach provides an upper bound estimate on the influence of age and sex and a conservative estimate on the influence of other measures, as all shared variance is attributed to age and sex. As shown in Fig. S6, the influence of body phenotypes dropped dramatically after removing shared variance with age and sex, indicating that their importance is primarily attributed to the shared variance.

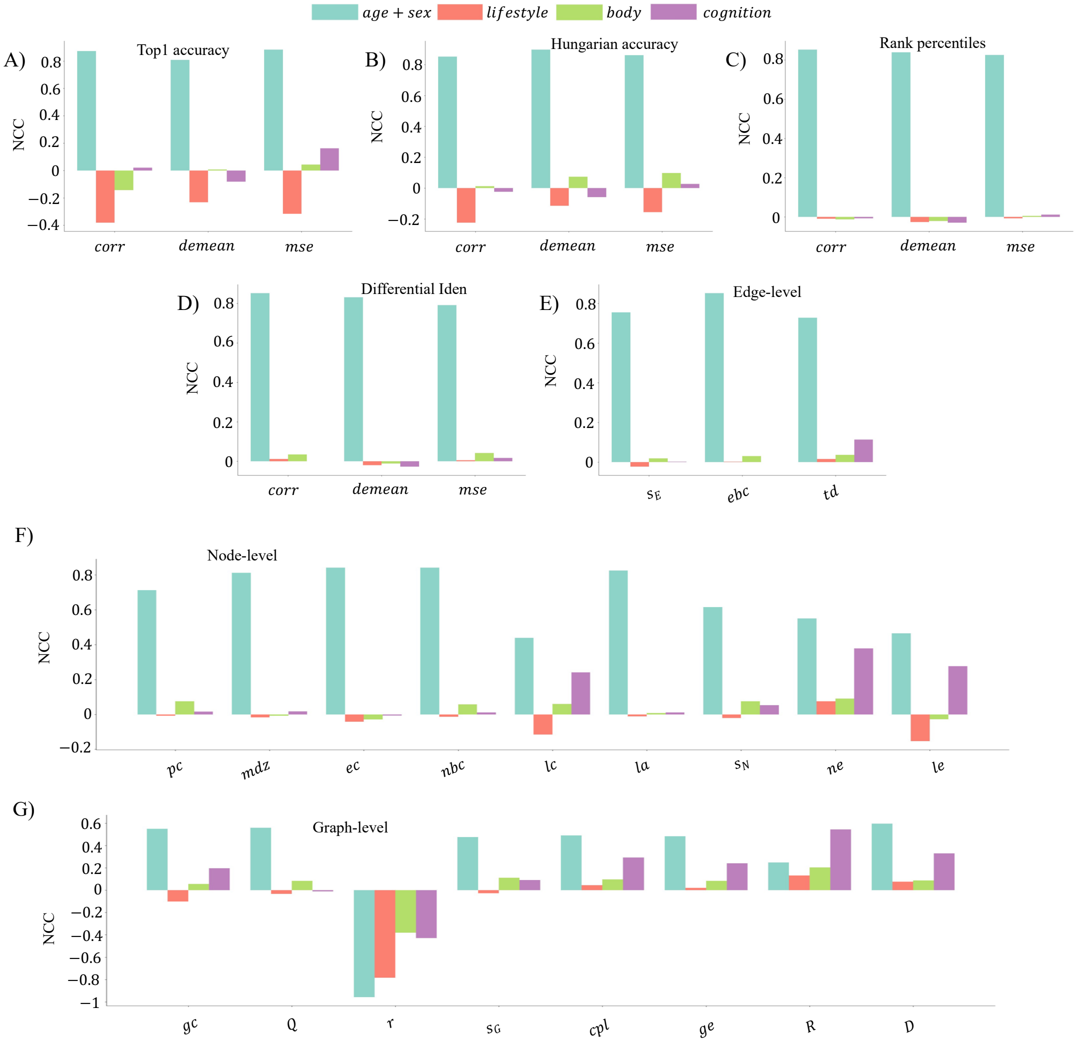


*Figure S6. NCC when age and sex were regressed out from other measures.*

#### ***Replicating generated variability in Desikan-Killiany atlas***

We replicated the results of generated variability, and different categories' contributions to variability generation, in the DK atlas consisting of 68 cortical regions. Here, Fig S7 quantifies the generated individual variation, while Fig S8 details each category's contribution to generating variability, in the DK atlas.

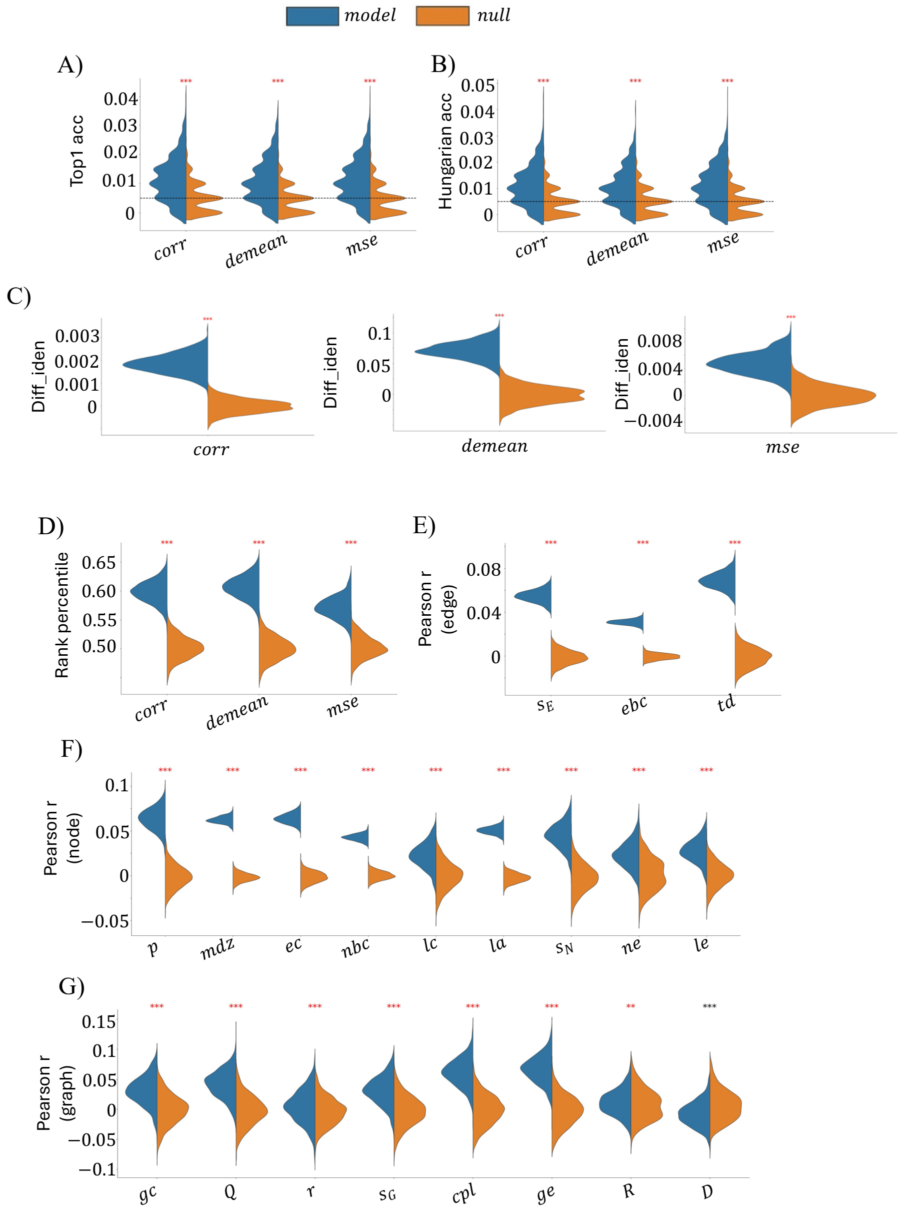


*Figure S7. Model outperforms null in generating individual variations. Same as Fig. S2 but in DK atlas.*

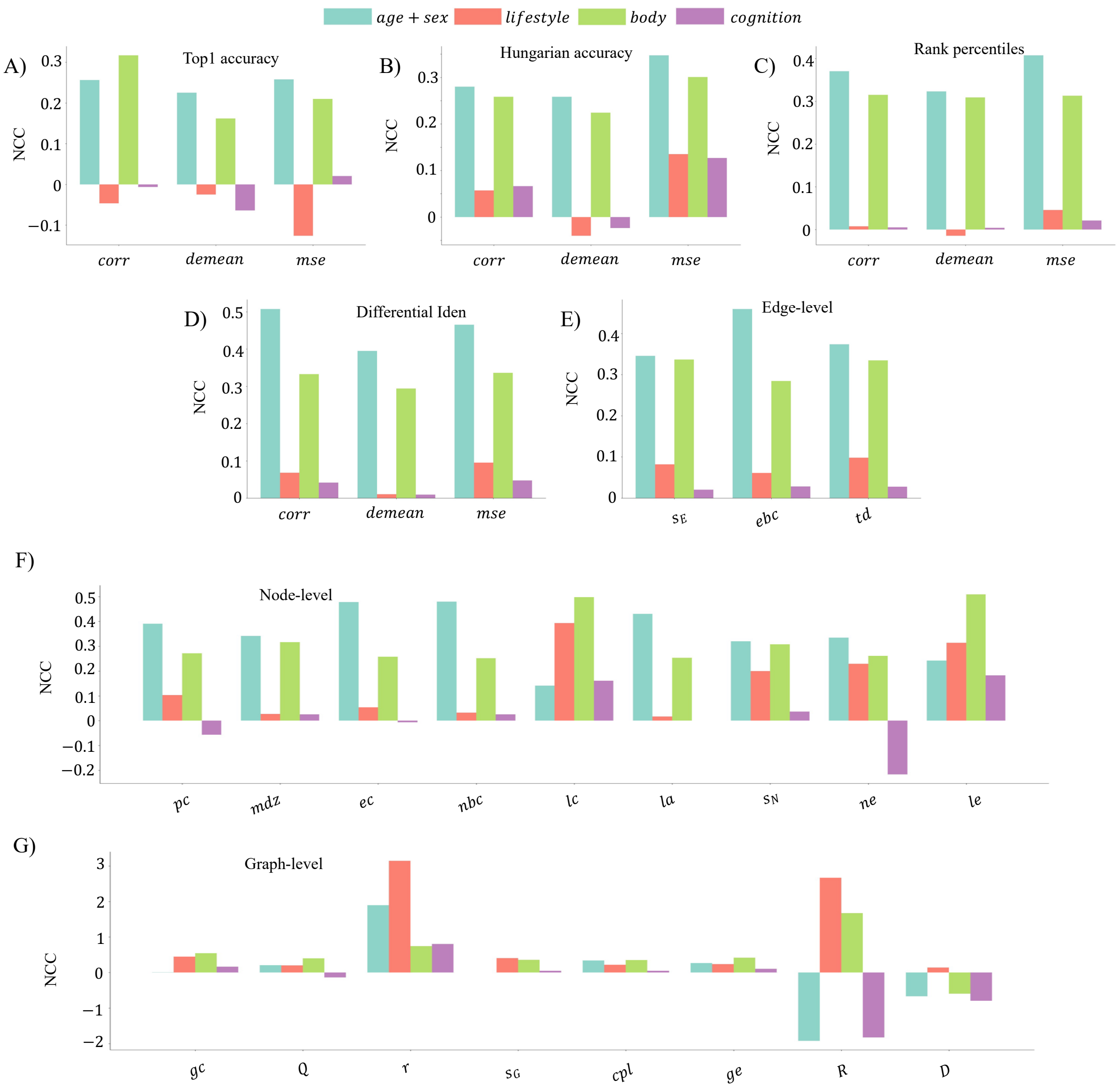


*Figure S8. NCC for all measures. Same as Fig. S4 but in DK atlas.*

##### *Intrinsic interindividual variation differences between brain subnetworks*

It is unclear if the differences in generated variation between brain subnetworks root in the fact that interindividual variations can intrinsically differ between regions in empirical brains. To investigate this, we employed connectomes mapped from the Human Connectome Project (Van Essen et al., 2013) test-retest scans because retest data are not available in UKB. Specifically, we computed the variation in test connectomes explained by the retest connectomes. Next, we evaluated whether the variation distribution among subnetworks was significantly different between model-generated and retest connectomes using a paired permutation test. A significant result suggests model-generated differences between subnetworks cannot be explained by the differences in empirical connectomes. We found patterns in empirical test-retest data (Fig. S9) are significantly different from model observations, suggesting that the heterogeneity in generated connectomes cannot be explained by the intrinsic distribution of interindividual variations.

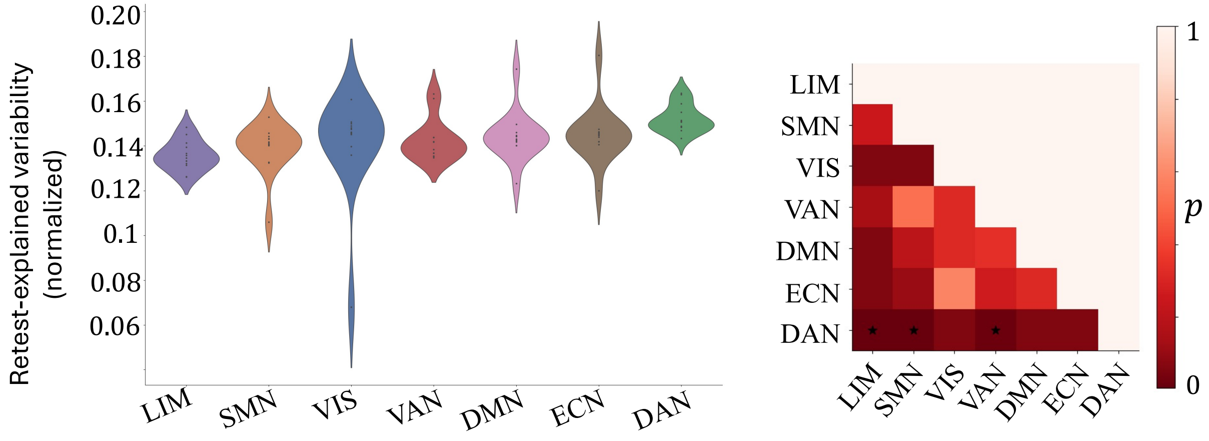


*Figure S9. Intrinsic interindividual variation distribution in empirical brains. Same as Figure. 4A in the main text but for test-retest captured variation.*

##### *Test for between-subnetwork differences in categorical contributions*

To assess whether the relative influence of personal data categories differed between subnetworks, we first computed each category's contribution (i.e., NCC) within each subnetwork. As shown in Fig. S10, the categorical contributions in a subnetwork were represented in a matrix (e.g., Matrix A of shape 13x4), where each row corresponded to an identifiability or network measure, and each column represented the NCC of a specific category within the subnetwork. For example, if row 1 of Matrix A represents differential identifiability, then A(1,2) quantifies the influence of lifestyle factors on generating variability measured by differential identifiability, within subnetwork 1.

To compare subnetworks, let matrices A and B represent the categorical contributions for subnetworks 1 and 2, respectively. We computed the Euclidean distance between row vectors of A and B, forming a distance matrix C of shape (26,26), where larger distances indicate dissimilar vectors. Elements enclosed within the triangular regions of C correspond to within-subnetwork distances ($\mathrm{D}_{\mathrm{within}-1}$ *and* $\mathrm{D}_{\mathrm{within}-2}$), and elements in the rectangular region represent between-subnetwork distances ($\mathrm{D}_{\mathrm{between}}$). The difference in the mean values of $\mathrm{D}_{\mathrm{between}}$ and $\mathrm{D}_{\mathrm{within}}$ quantifies how dissimilar the categorical contributions were between subnetworks 1 and 2.

To quantify the significance of the dissimilarity, we performed paired permutation test. In each permutation, a subset of rows was randomly selected and swapped between matrices A and B, forming permuted matrices C and D. For the example in Fig. S10, rows highlighted with black rectangles in matrices C and D were swapped. We computed the differences between the mean values of $D_{between}$ and $D_{within}$ (matrix E) using the permuted matrices. This process was repeated for 10,000 times to generate a null distribution of dissimilarity. The observed dissimilarity was then evaluated against this null distribution to determine statistical significance.

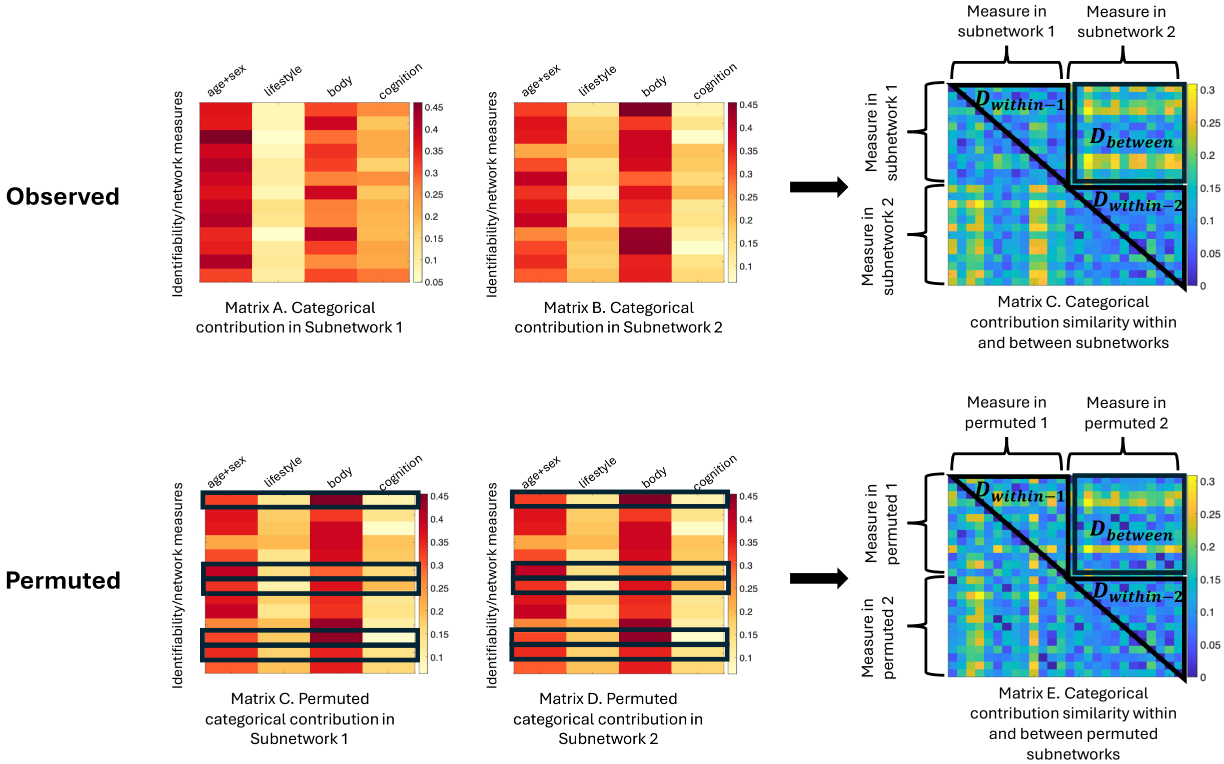


*Figure S10. Schematic illustration on the process that compares categorical contributions between subnetworks.*

#### References


Akarca, D., Vértes, P. E., Bullmore, E. T., & Astle, D. E. (2021). A generative network model of neurodevelopmental diversity in structural brain organization. *Nature communications, 12*(1), 1-18.

Betzel, R. F., Avena-Koenigsberger, A., Goñi, J., He, Y., De Reus, M. A., Griffa, A., Vértes, P. E., Mišic, B., Thiran, J.-P., & Hagmann, P. (2016). Generative models of the human connectome. *Neuroimage, 124*, 1054-1064.

Jamison, K. W., Gu, Z., Wang, Q., Sabuncu, M. R., & Kuceyeski, A. (2024). Release the Krakencoder: A unified brain connectome translation and fusion tool. *bioRxiv*.

Liu, Y., Seguin, C., Mansour, S., Oldham, S., Betzel, R., Di Biase, M. A., & Zalesky, A. (2023). Parameter estimation for connectome generative models: Accuracy, reliability, and a fast parameter fitting method. *Neuroimage, 270*, 119962.

Oldham, S., Fulcher, B. D., Aquino, K. M., Arnatkeviciute, A. M., Paquola, C., Shishegar, R., & Fornito, A. (2021). Modeling spatial, developmental, physiological, and topological constraints on human brain connectivity. *bioRxiv*.

Van Essen, D. C., Smith, S. M., Barch, D. M., Behrens, T. E., Yacoub, E., Ugurbil, K., & Consortium, W.-M. H. (2013). The WU-Minn human connectome project: an overview. *Neuroimage, 80*, 62-79.

Zhang, X., Braun, U., Harneit, A., Zang, Z., Geiger, L. S., Betzel, R. F., Chen, J., Schweiger, J. I., Schwarz, K., & Reinwald, J. R. (2021). Generative network models of altered structural brain connectivity in schizophrenia. *Neuroimage, 225*, 117510.
